# Synaptic sign switching mediates online dopamine updates

**DOI:** 10.1101/2025.07.23.666367

**Authors:** Shun Li, Wengang Wang, Grace Knipe, Elliot Jerng, Paolo Capelli, Catherine Zhou, Eliana Bilsel, Bernardo L. Sabatini

## Abstract

In the mammalian brain, excitatory and inhibitory synapses are generally distinct and have fixed synaptic signs. Therefore, unlike in artificial neural networks, learning in biological networks is thought to be manifested by plasticity mechanisms that modify synaptic weights but not signs. Here, we demonstrate experience-dependent sign switching at synapses between glutamate and GABA co-releasing neurons of the entopedunculus (EP) and their targets in the lateral habenula (LHb). Pairing of reward or punishment with activation of EP co-releasing neurons makes EP-LHb synapses relatively more inhibitory or excitatory, respectively. Synaptic sign switching modulates downstream dopaminergic signaling, correlates with recent dopamine updates, and contributes to reinforcement learning. These data unveil a plasticity mechanism that alters both synaptic signs and weights to rapidly update dopamine release and drive learning.

## Main text

In most of the mammalian brain, the excitatory neurotransmitter glutamate and the inhibitory neurotransmitter GABA are released at pre-synaptic terminals of different neurons. Therefore, the sign of each synapse is typically fixed as excitatory or inhibitory, defined in terms of the ionic currents evoked by neurotransmitter release in its postsynaptic partner. For this reason, experience- and activity-dependent synaptic plasticity mechanisms are typically described as altering synaptic strengths but not synaptic signs (*1, 2*). This feature of biological neural networks is in strong contrast with the structure of artificial neural networks (ANNs) in which each connection (the equivalent of a synapse) between two nodes (neuron equivalents) can be either positive or negative and can change during training. Sign-switching plasticity increases performance in ANN by expanding the solution space of a network (*2, 3*).

However, the mammalian brain does contain synapses that are at least theoretically possible to implement sign-switching plasticity. For example, at synapses where glutamate and GABA are released by the same presynaptic bouton, the numbers of ionotropic glutamate and GABA receptors in the apposed post-synaptic terminal could determine both the synaptic sign and strength. One group of candidate sign-switching synapses are those formed between somatostatin (Sst)-expressing neurons in the entopeduncular nucleus (EP) and their targets in the lateral habenula (LHb). EP Sst neurons co-package glutamate and GABA into individual synaptic vesicles such that each vesicle exocytosis event co-releases these opposing neurotransmitters onto LHb neurons (*4, 5*). Furthermore, individual EP Sst→LHb synapses show both structural and functional evidence of co-transmission of glutamate and GABA, indicating that individual postsynaptic terminals contain receptors for both transmitters (*5*). Nevertheless, postsynaptic currents in LHb activated by release of individual vesicles from EP Sst axons exhibit a wide range of glutamatergic and GABAergic current amplitudes, indicating inter-synapse variability in both synaptic sign and strength (*5*).

The circuitry of the EP and LHb suggests a specific function for these glutamate/GABA co-releasing synapses. EP Sst neurons project exclusively to LHb (*4*), signal information about movements, goals, and value, which crucially contributes to valence encoding in LHb (*6–8*). The LHb is a central component of a neural circuit that is responsible for calculating the expected value of specific contexts and actions: glutamatergic neurons in the LHb activate GABAergic neurons in the rostromedial tegmental nucleus (RMTg) and ventral tegmental area (VTA), which di-synaptically inhibit dopaminergic neurons of the VTA and thereby lower dopamine (DA) levels in the nucleus accumbens (NAc) (*9–12*). Experiences shape LHb activity such that it typically signals negative value or negative value prediction, contributing to the calculation of reward prediction error (RPE) that is encoded by VTA DA neurons (*12–17*). Curiously, LHb are enriched with inputs from neurons capable of co-releasing both glutamate and GABA (*4, 18–22*). Therefore, sign-switching plasticity at inputs to the LHb, if present, might enable rapid updates of value and RPE calculation, consistent with the function of LHb in online adaptive learning (*23–31*).

Here, we propose that EP Sst→LHb synapses undergo experience-dependent sign-switching plasticity that alters their relative glutamatergic and GABAergic transmission. Furthermore, we propose that sign-switching plasticity in these synapses mediates the rapid updating of downstream RPE-encoding DA transients in the NAc. We test this hypothesis using a novel behavioral paradigm in which the valence of EP Sst neuron activity is rapidly updated by experience, triggering plasticity at the EP-LHb-VTA circuit. *In vivo* photometric analysis of DA release and other neural signals followed by *ex vivo* analysis of synaptic transmission support the existence of experience-dependent synaptic sign switching plasticity at EP Sst→LHb synapses that modulates DA release in NAc and supports fast adaptive learning.

## Results

We hypothesize that experience-dependent plasticity determines the relative contribution of glutamatergic and GABAergic transmission at synapses between EP Sst neurons and their targets in the LHb. Furthermore, we propose that this plasticity rapidly modulates downstream DA signaling as environmental contingencies change (**Fig. 1A**). Examining this hypothesis requires protocols to: (1) induce plasticity in a cell type-specific and experience-dependent manner; (2) measure the net effect of synaptic activity on value computation and behavior *in vivo*; and (3) measure the glutamate vs. GABA transmission ratio of behaviorally relevant synapses *ex vivo*.

**Figure 1.**
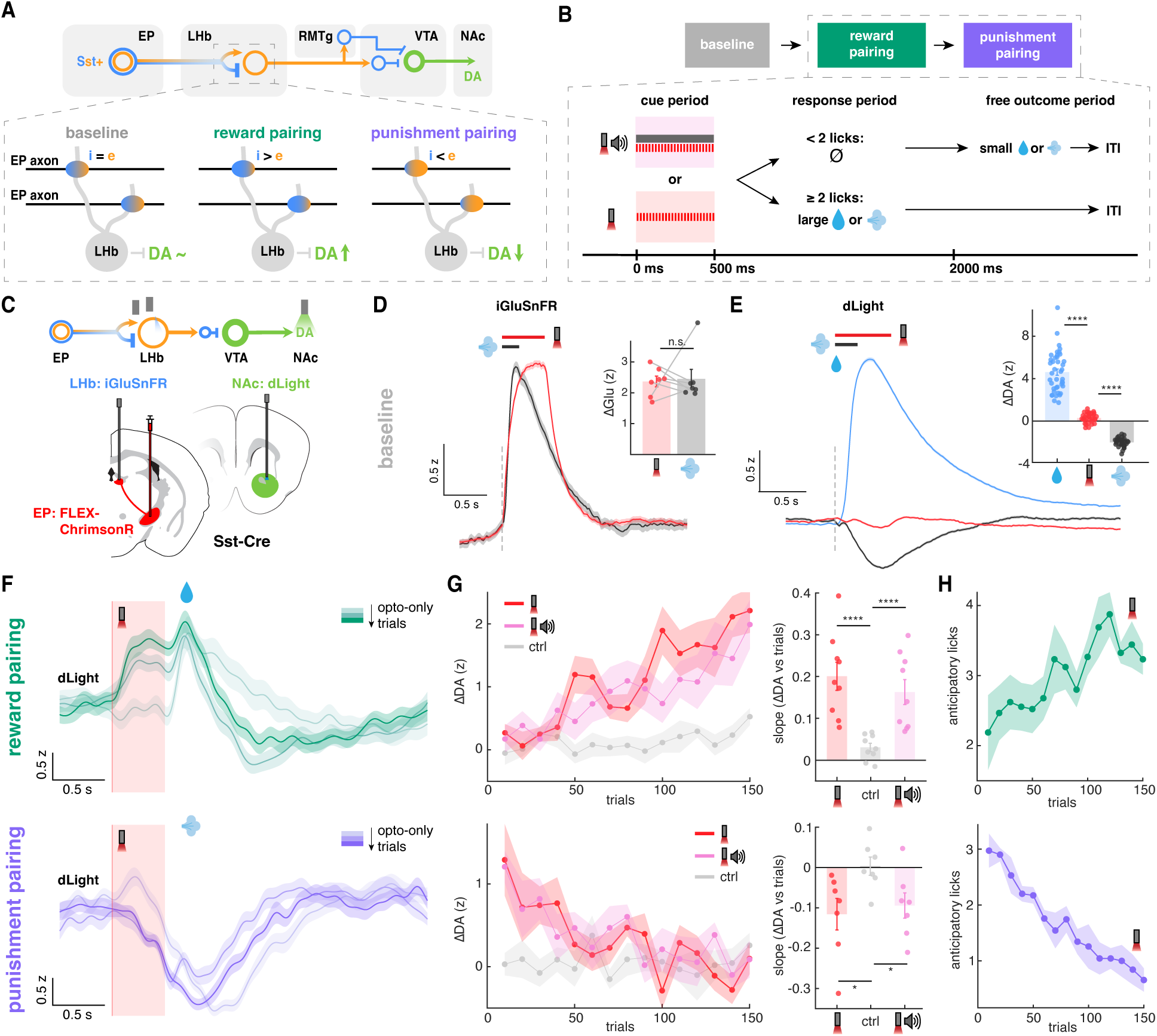
EP Sst glutamate & GABA co-releasing neurons bi-directionally modulate dopamine through opto-outcome pairing. **A**, Schematics of the EP-LHb-VTA circuit that regulates NAc DA signaling (*top*) and the proposed model of sign-switching synaptic plasticity (*bottom*). *top*, EP Sst neurons project exclusively to LHb where they co-release glutamate and GABA. LHb neurons synapses onto GABAergic neurons in RMTg or VTA to disynaptically inhibit NAc-projecting DA neurons. *bottom*, We hypothesize that, at baseline, excitatory and inhibitory synaptic currents at EP-LHb synapses roughly cancel out and resulting in little or modulation of NAc DA. However, if activity the EP axons correlates with reward (*middle*), plasticity at EP-LHb synapses increases the relative strength of inhibitory transmission, resulted in inhibition of LHb and disinhibition of DA release. Conversely, if activity of EP axons correlates with aversive outcomes (punishment, *right*), plasticity at EP-LHb synapses increases the relative strength of excitatory transmission, resulted in excitation of LHb and inhibition of DA release. **B**, Overview of opto-pairing paradigm used to test the model. *top*, Each mouse experiences three behavioral paradigms during which NAc dopamine and other brain signals are monitored. The “baseline” phase consists of sessions in which different stimuli (water reward, air puff to the eye, tones, and optogenetic stimulation of EP Sst neurons) are given in random order with no pairing. In the “reward pairing” or “punishment pairing” phases, optogenetic activation of EP Sst neuron axons in the LHb (referred as “EP opto stim”) is positively or negatively reinforced by consistent pairing with rewarding (water) or aversive (air puff) outcomes, respectively. *bottom*, During the reinforcement blocks, most randomly selected cues that consist of either (1) 500 ms of simultaneous EP opto stim and a continuous tone or (2) 500 ms of EP opto stim alone are presented. If in the following response period, the animal licks 2 or more times, they immediately receive either a large reward (for reward pairing sessions) or large air puff to the eye (for punishment pairing sessions). If they do not lick at least two times, a small reward or air puff is delivered at the start of the free outcome period. A random length inter-trial interval (ITI) occurs before the next trial. **C**, Schematic of viral transductions and photometric recordings. For all animals, Cre-dependent ChrimsonR (FLEX-ChrimsonR) is selectively expressed in EP Sst neurons in *Sst-Cre* mice. An optical fiber is implanted in LHb to optogenetically stimulate ChrimsonR+ axons of EP Sst neurons. The fluorescent dopamine sensor dLight is expressed in NAc to monitor DA release through a second optical fiber. In a subset of animals, fluorescent glutamate sensor iGluSnFR is expressed in LHb so that glutamate release can be monitored via photometry through the same optical fiber used for optogenetic stimulation. **D,** Glutamate transients in LHb as reported by iGluSnFR fluorescence increases in response to either an air puff (black) or the EP opto stim (red) during the baseline phase. The averaged z-scored signal ± s.e.m. is shown. Inset, animal-averaged amplitudes of the evoked glutamate transients in LHb (ΔGlu) (n=7 mice). **E**, NAc DA transients during the baseline phase as reported by dLight fluorescence aligned to onset of water delivery (blue), the EP opto stim (red), and air puff (black). The average z-scored signal ± s.e.m. is shown. Inset, animal-averaged amplitudes of the evoked NAc DA transients (ΔDA) (water: n=41 mice; EP opto stim: n=33 mice; air puff: n=33 mice) **F**, NAc DA transients aligned to onset of opto-only trials during reward (*top*) and punishment (*bottom*) pairing sessions for an example animal. Trials are color-coded from light to darker color indicating early to late trials (each color is 30 opto-only trials). The average z-scored signals ± s.e.m. are shown. The icons indicate the approximate times of water or air puff delivery. **G**, The amplitude of cue-evoked NAc DA transients (ΔDA) as a function of trial number (*left*) pairing and the corresponding slope between changes in evoked DA transients and trial number (*right*, ΔDA vs trials). Reward pairing data are plotted on the top (n=9 mice) and punishment data are plotted at the bottom (n=7 mice). Data are shown for trials with the opto-only cue (red), opto+tone cue (pink), or randomly selected recording timepoints throughout each session as control (gray). On the left, each dot indicates the average DA amplitudes of the nearby 10 trials and the averaged z-scored amplitudes across animal ± s.e.m. are shown. On the right, the data for individual mice (dots) and the average ± s.e.m are shown. **H,** The number of anticipatory licks (i.e., those that occur during the 500 ms cue period) for opto-only cue trials in reward (*top*, n=9 mice) and punishment (*bottom,* n=7 mice) pairing sessions as a function of trial number (binned as in panel **G**). The average number of anticipatory licks across animal ± s.e.m. is shown.

To achieve the first two goals, we designed an optogenetic paradigm to rapidly update the valence associated with the activity of subsets of EP Sst glutamate/GABA co-releasing neurons while continually monitoring the effect of the EP optogenetic stimulus on the EP-LHb-VTA circuit as reflected by DA release in NAc (**Fig. 1A, B**). We achieved this by optogenetic stimulating EP co-releasing neurons by expressing the excitatory opsin ChrimsonR in EP of *Sst-ires-Cre* mice. In all behavioral sessions, we performed continuous DA measurements at NAc with fiber photometry using the DA sensor dLight (**Fig. 1C**; **fig. S1**, **S2**). Thirsty mice were trained to either lick or withhold licking following optogenetic activation of EP Sst neurons (referred as “EP opto stim”) by operantly delivering either a water reward or air puff to the eye triggered by licking. In sessions in which EP opto stim was associated with water delivery, the mouse was motivated to lick following the stimulus presentation to receive the positive reinforcer. In contrast, in sessions in which EP opto stim was associated with air puffs, the mouse was motivated to withhold licking to avoid the negative reinforcer. The pairing of EP opto stim with reward or punishment varied across behavioral sessions (and constant within each session), such that animals needed to learn the association and rapidly induce plasticity (**Fig. 1B**).

We predicted that pairing EP opto stim with positive outcomes (water) would result in it gaining the ability to increase NAc DA by rendering EP Sst→LHb synapses more GABAergic. On the other hand, subsequent pairing with negative outcomes (air puff) would reverse this association, rapidly decreasing the amplitude of EP opto stim evoked transients in NAc DA and rendering EP Sst→LHb synapses more glutamatergic (**Fig. 1A**). Furthermore, we predicted that these changes are mirrored by behavioral changes: pairing with a positive reinforcer results in EP opto stim driving licking, which is rapidly reversed by pairing with a negative reinforcer.

## EP Sst glutamate and GABA co-releasing neurons bi-directionally modulate DA

Water-restricted mice first experienced baseline sessions in which EP opto stim, water delivery, air puffs, and tones were delivered at random times. These stimuli are delivered without any contingency between them, allowing us to assess the baseline state of the EP-LHb-VTA circuit. We calibrated the power and pattern of the EP opto stim so that the peak level of glutamate release it evoked in the LHb (as measured by photometric recording of glutamate sensor iGluSnFR) is similar to that evoked by an aversive air puff (**Fig. 1D**). NAc DA reliably increases and decreases in response to delivery of water and air puffs, consistent with the stimuli being intrinsically associated with reward and punishment, respectively (**Fig. 1E**). However, EP opto stim during baseline sessions does not modulate NAc DA significantly, despite triggering substantial glutamate release in LHb (**Fig. 1D, E**). This suggests that activity of EP Sst→LHb synapses, as a population, has neutral valence at baseline, consistent with the glutamatergic and GABAergic signaling largely canceling out and having little effect on downstream DA activity.

We next paired EP opto stim with water and then with air puffs across different sessions (**Fig. 1F-H**). Two main types of cues are delivered in approximately equal portion during each session: one consists of EP opto stim alone (“opto-only trials”, shown in red in all figures) and the other of simultaneous presentation of EP opto stim and a 3 kHz auditory tone (“opto+tone trials”, shown in pink in all figures) (**movie S1**). Both cues last 500 ms, which is long enough to examine NAc DA transients and anticipatory licking during the cue period before the reinforcer is delivered. The opto+tone trials help the animals learn the EP opto stim contingency and serve as an important positive control for learning (see next section). Trials of each type are randomly interleaved. Animals reliably learned the opto-outcome association such that both the opto-only and opto+tone trials gained the ability to manipulate NAc DA and behavior (**Fig. 1G, H**; **fig. S3A, C**; **movie S2**). Specifically, during reward pairing, EP opto stim gradually gains the ability to induce positive DA transients in NAc (**Fig. 1F, G**; **fig. S3C**) as well as anticipatory licking during the cue presentation (**Fig. 1H**; **fig. S3A, C**; **movie S2**). Thus, pairing optogenetic activation of EP Sst inputs to LHb with rewards changes the output of EP-LHb-VTA circuit, resulting in NAc DA signals that are consistent with the formation of cue-outcome associations by positive reinforcement.

Following reward pairing, we reversed the opto-outcome contingency by pairing the EP opto stim and opto+tone cues with air puffs. Because this change was not explicitly indicated to the animal, at the beginning of the session EP opto stim still induced positive DA transients and anticipatory licking during the cue period (**Fig. 1F-H**; **fig. S3B, D**; **movie S2**). However, repeated pairing decreased and the eventually eliminated both positive DA transients in response to EP opto stim and anticipatory licking during the cue period (**Fig. 1F-H**; **fig. S3B, D**; **movie S2**).

These data indicate that the initially neutral EP Sst neurons gain the ability to positively modulate DA release in NAc through pairing of EP opto stim with reward. This effect can also be rapidly reversed by changing environmental conditions, in our case by pairing with a negative reinforcer.

Several controls discounted the possibility that the animal guides its behavior using the laser light or other sensory cues rather than the activation of EP Sst neurons (for details see Methods section “*Behavior: preventing mice from cheating*”). First, we injected Cre-dependent viruses into *Sst-ires-Flp* or WT mice, thereby preventing opsin expression in EP (**fig. S4A, B**). These mice failed to learn pairings of opto-only cue to outcome, evidenced by the lack of changes in NAc transients and anticipatory licking in response to opto-only cues across reward and punish pairing trials (**fig. S4D, E**). In the same sessions, the animals did learn the opto+tone-outcome pairing due to the presence of tone, indicating that learning of opto-only trials specifically requires optogenetic activation of EP neurons (**fig. S4D, E**). Second, we expressed ChrimsonR in ventral thalamic reticular nucleus (TRN), which is near the EP, contains Sst-expressing neurons, and can be accidently transduced with AAV in *Sst-Cre* mice (**fig. S5A-C**). *Ex vivo* whole-cell recordings in acute brain slices revealed a lack of monosynaptic connections between ventral TRN and LHb neurons (0/21 cells, N=4 mice, **fig. S5D, E**). Consistent with this, animals with ChrimsonR expressed in TRN Sst neurons also did not learn the opto-outcome association (**fig. S5F, G**). This result also demonstrates that not all optogenetic stimuli can drive behavior and NAc DA transients via positive reinforcement.

Together, our data demonstrate that EP Sst co-releasing neurons in the baseline state are neutral such that their activity does not modulate NAc DA, but they gain the ability to bi-directionally modulate of NAc DA once their activity is paired with rewarding or aversive outcomes.

## Bi-directional modulation of DA by activity of EP Sst neurons is LHb dependent

EP Sst neurons project only to LHb which, in turn, di-synaptically inhibits VTA DA neurons (*4, 10*). We find that effects of EP Sst neurons on NAc DA release and behavior are modulated by positive and negative reinforcers, suggesting that plasticity occurs in the EP-LHb-VTA circuit. To narrow the possible locus that mediates this circuit plasticity, we performed four groups of experiments.

First, if the reinforcement-induced changes of EP activity on NAc DA depend on LHb, then lesioning LHb neurons should abolish learning. We unilaterally injected AAV encoding Cre-dependent caspase into LHb of *Vglut2-Cre* mice, effectively ablating most LHb neurons (**Fig. 2A**; **fig. S6A, B**). Animals with LHb lesioned in the same hemisphere in which the optogenetic activation was delivered failed to learn opto-outcome pairing, showing no consistent NAc DA transients (**Fig. 2B, C**) and anticipatory licking (**fig. S6D**) in response in opto-only trials. Several lines of evidence collectively suggest that deficit in learning opto-outcome pairing is specific to caspase ablation. Firstly, water rewards and aversive air puffs still increased and decreased NAc DA in these mice, respectively, confirming they have the intact ability to detect both stimuli at their innate valence as well as our continued ability to detect DA transients (**fig. S6C**).

**Figure 2.**
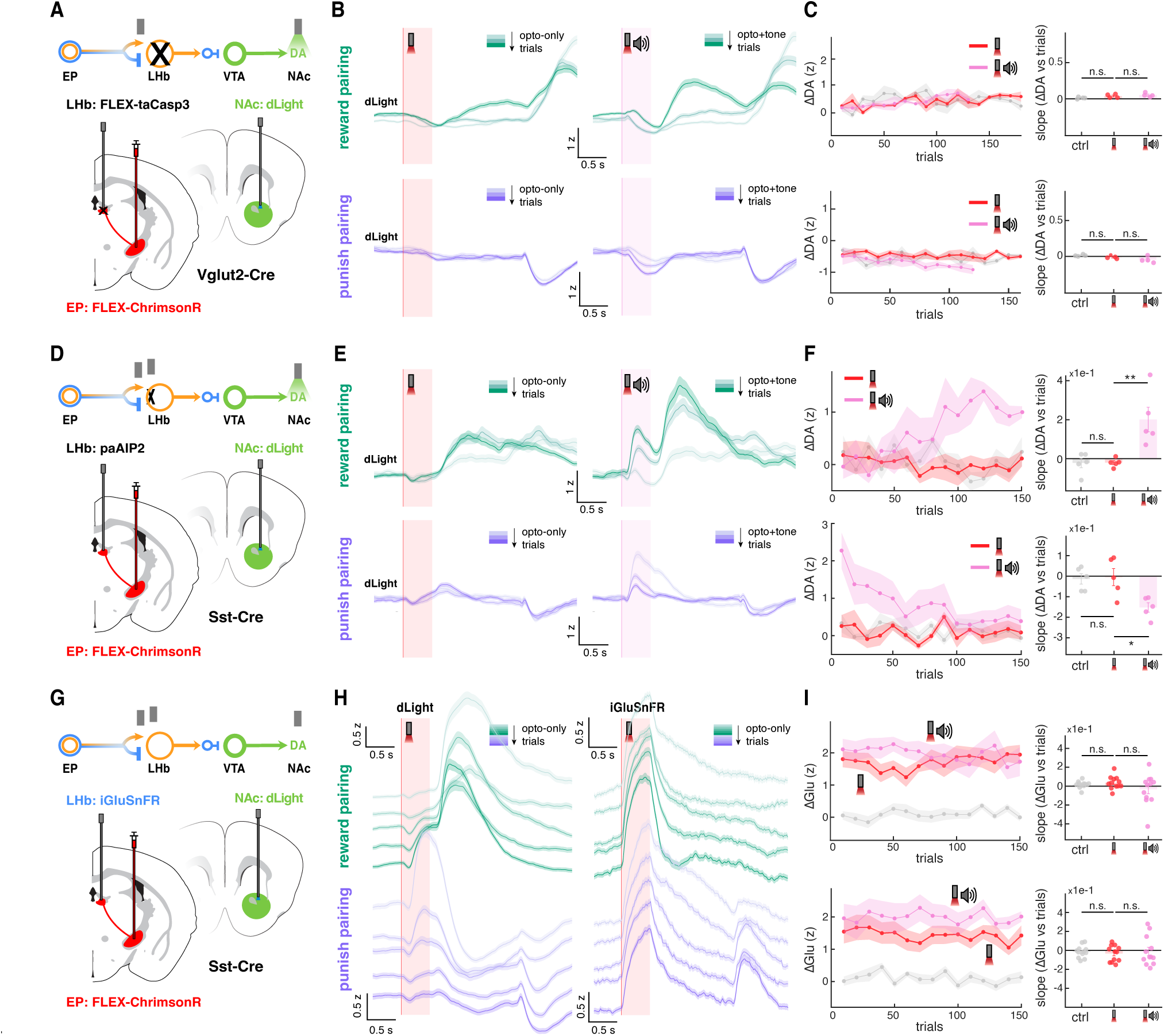
Bi-directional modulation of dopamine by EP is LHb dependent **A**, Schematic of viral transductions and photometric recordings for LHb taCasp3 experiment. Cre-dependent ChrimsonR and Cre-dependent taCasp3 are selectively expressed in EP and LHb of the *Vglut2-Cre* mice. An optical fiber is implanted in LHb to optogenetically stimulate EP ChrimsonR+ axons in LHb. dLight is expressed in NAc to monitor DA release through a second optical fiber. **B**, NAc DA transients aligned to onset of opto-only cue (left, red) or opto+tone cue (right, pink) during reward pairing sessions (*top*) and punishment pairing sessions (*bottom*) (n=4 mice). Trials are color-coded from light to darker color indicating early to late trials (each color is around 200 trials across all animals, i.e. 50 trials for each animal). The average z-scored signals across trials from all animals ± s.e.m. are shown. **C**, The amplitude of cue-evoked NAc DA transients (ΔDA) as a function of trial number (*left*) and corresponding slope between changes in evoked DA transients and trial number (*right*, ΔDA vs trials). Reward pairing data are plotted on the top and punishment pairing are plotted at the bottom. Data are shown for trials with the opto-only cue (red), opto+tone cue (pink), or randomly selected recording timepoints throughout each session as control (gray). On the left, each dot indicates the average DA amplitudes of the nearby 10 trials and the averaged z-scored amplitudes across animal ± s.e.m. are shown. On the right, the data for individual mice (dots) and the average ± s.e.m are shown. **D**, Schematic of viral transductions and photometric recordings for LHb paAIP2 experiment. Cre-dependent ChrimsonR is selectively expressed in EP Sst neurons and paAIP2 is expressed in LHb of *Sst-Cre* mice (n=5 mice). An optical fiber is implanted in LHb to optogenetically stimulate EP ChrimsonR+ axons in LHb and deliver periodic blue light to induce paAIP2-mediated inhibition of CaMKII throughout the session. Fluorescent dopamine sensor dLight is expressed in NAc to monitor DA release through a second optical fiber. **E**, NAc DA transients aligned to onset of opto-only cue (left, red) or opto+tone cue (right, pink) during reward pairing sessions (*top*) and punishment pairing sessions (*bottom*). Trials are color-coded from light to darker color indicating early to late trials (each color is around 250 trials across all animals, i.e. 50 trials for each animal). The average z-scored signals across trials from all animals ± s.e.m. are shown. **F**, The amplitude of cue-evoked NAc DA transients (ΔDA) as a function of trial number (*left*) and the corresponding slope between changes in evoked DA transients and trial number (*right*, ΔDA vs trials). Reward pairing data are plotted on the top and punishment pairing data are plotted at the bottom. Data are shown for trials with the opto-only cue (red), opto+tone cue (pink), or randomly selected recording timepoints throughout each session as control (gray). On the left, each dot indicates the average DA amplitudes of the nearby 10 trials and the averaged z-scored amplitudes across animal ± s.e.m. are shown. On the right, the data for individual mice (dots) and the average ± s.e.m are shown. **G**, Schematic of viral transductions and photometric recordings for LHb iGluSnFR experiment. For all animals (n=12 mice), Cre-dependent ChrimsonR is selectively expressed in EP Sst neurons of *Sst-Cre* mice. Fluorescent glutamate sensor iGluSnFR is expressed in LHb to monitor local glutamate transients in LHb. An optical fiber is implanted in LHb to optogenetically stimulate EP ChrimsonR+ axons in LHb and perform fiber photometry for iGluSnFR. In 9 out of 12 animals, fluorescent dopamine sensor dLight is expressed in NAc to monitor DA release through a second optical fiber. **H**, NAc DA transients (*left*) and LHb glutamate transients (*right*) aligned to onset of opto-only cue (left, red) or opto+tone cue (right, pink) during reward pairing sessions (*green*) and punishment pairing sessions (*purple*). Trials are color-coded from light to darker color indicating early to late trials (each color is the average of nearby 30 trials from all animals). The average z-scored signals across trials from all animals ± s.e.m. are shown. **I**, The amplitude of cue-evoked LHb glutamate transients (ΔGlu) as a function of trial number (*left*) and the corresponding slope between changes in evoked Glu transients and trial number (*right*, ΔGlu vs trials). Reward pairing data are plotted on the top and punishment pairing data are plotted at the bottom. Data are shown for trials with the opto-only cue (red), opto+tone cue (pink), or randomly selected recording timepoints throughout each session as control (gray). On the left, each dot indicates the average Glu amplitudes of the nearby 10 trials and the averaged z-scored amplitudes across animal ± s.e.m. are shown. On the right, the data for individual mice (dots) and the average ± s.e.m are shown.

Furthermore, LHb ablated mice still successfully learn and reverse tone-outcome pairings when the tone-only cue is presented >80% of trials (**fig. S6E, F**), ruling out gross cognitive deficits due to unilateral LHb lesion. Since opto+tone cues are presented intermixed with opto-only cues during original pairing sessions (**Fig. 2B, C**), LHb lesions likely render opto-only cues undetectable by downstream circuits, essentially turning these trials into “free rewards”. As these are ∼50% of trials, this likely leads to contingency degradation that abolishes learning in opto+tone trials as well. Lastly, *Vglut2-Cre* animals without LHb lesions can successfully learn and reverse the opto-outcome pairing (**fig. S7**), confirming that the observed deficits are specific to caspase ablation and do not arise from the stimulation of EP Vglut2→LHb synapses, which are formed by both Sst neurons and the purely glutamatergic PV neurons (also solely project to LHb) (*4*).

Second, if postsynaptic plasticity of LHb neurons is blocked or impaired, animals should not learn the opto-outcome pairing. Previous studies have showed that CaMKII is a key mediator of LHb plasticity (*32*). We therefore unilaterally expressed photo-activatable CaMKII inhibitor paAIP2 (*33*) in LHb, ChrimsonR in EP Sst→LHb neurons, and dLight in NAc (**Fig. 2D**; **fig. S6A, B**). During opto-pairing sessions, paAIP2 was turned on and prevented the emergence of optogenetically evoked DA transients during the cue period of opto-only trials (*red*, **Fig. 2E, F**). Anticipatory licking behavior in response to the opto-only cue was also abolished (**fig. S6H**).

Moreover, this effect was not due to gross cognitive deficits as animals successfully acquired tone-outcome pairing in opto+tone trials (*pink*, **Fig. 2E, F**) and all animals later learned opto-reward pairing when CaMKII was not inhibited (**fig. S6I, J**). Together with the LHb lesion experiment, these manipulations indicate that LHb neurons and CaMKII dependent signaling are necessary for the activity of EP Sst neurons to gain the ability to modulate DA during opto-pairing task.

Third, to investigate potential presynaptic contributions to the circuit plasticity, we monitored neurotransmitter release in LHb during learning. Unfortunately, currently available fluorescence GABA sensors lack the sensitivity necessary to record GABA release in LHb during training.

However, we were able to record glutamate transients in LHb using iGluSnFR and DA in NAc throughout all sessions of the opto-pairing task (**Fig. 2G**). If EP-LHb-VTA plasticity were primarily driven by presynaptic changes in the ratio of glutamate versus GABA co-packaged and released by EP Sst neurons, we would expect clear shifts in glutamate transients across different phases of pairing. All animals successfully acquired the opto-outcome association, as reflected in EP opto stim-induced DA (**Fig. 2H**) and anticipatory licking (**fig. S3C, D**) increases during reward pairing and subsequent reversal of DA transients during punishment pairing. However, glutamate transients in LHb during cue periods remained stable throughout learning across both opto-only and opto+tone trials (**Fig. 2H, I**). Thus, glutamate release in LHb following activation of EP Sst axons is not affected by positive and negative reinforcement of EP opto stim.

Lastly, we moved downstream to test whether LHb activity maintains a fixed inhibitory influence on NAc DA release regardless of the direction of opto-outcome pairing. To examine this, we expressed a light-gated potassium channel WiChR (*34*) to transiently inhibit LHb neurons while recording NAc DA transients (**fig. S8A, B**). Consistent with previous literature (*24–27*), inhibition of LHb transiently increases DA in NAc (**fig. S8C**). This inhibitory effect of LHb activity on NAc DA was independent of the nature of EP opto stim pairing: WiChR-mediated LHb inhibition reliably induces DA increases of similar amplitude after both reward and punish pairing sessions (**fig. S8C, D**). Thus, LHb as a population consistently suppresses NAc DA throughout different phases of opto-outcome pairing, consistent with a fixed negative sign between LHb and DA release throughout learning.

Together, these experiments collectively support that bi-directional modulation of DA transients and behavior by EP opto stim is mediated by plasticity at EP Sst→LHb synapses. The dependence on postsynaptic CaMKII activity and the lack of change in glutamate release in LHb suggest that this plasticity is primarily driven by post-synaptic mechanisms.

## Sign switching occurs at EP Sst**→**LHb synapses during learning

To directly assess the sign of EP Sst→LHb synapses and its potential to switch upon different opto-outcome pairings, we performed *ex vivo* electrophysiology recordings after *in vivo* opto-pairing sessions. We exploited the optogenetic cue used to activate EP Sst neurons *in vivo* to allow us to target the axons of the same population in *ex vivo* brain slices. The optogenetic cue thus act as a bridge, permitting further examination of those same synapses *ex vivo* that underwent plasticity *in vivo*.

We performed *ex vivo* whole-cell electrophysiological recordings at various stages after the opto-pairing task (**fig. S9**). Specifically, we optogenetically stimulated terminals of EP Sst neurons in LHb and recorded excitatory and inhibitory postsynaptic currents (EPSCs and IPSCs) in voltage-clamped LHb neurons (held at -70 mV for EPSCs and 10 mV for IPSCs, corresponding to the effective reversal potentials measured empirically for each current without adjustment for liquid junction potentials) (**Fig. 3A**). To isolate monosynaptic inputs, we included TTX and 4-AP in all recordings. We also blocked NMDA-type glutamate receptors to isolate fast currents mediated by AMPA and GABA_A_ receptors. To quantify and summarize the relative contribution of excitatory and inhibitory currents, we calculated a “synaptic sign index” for each cell, which measures the difference over the sum of the amplitudes of IPSCs and EPSCs (see Methods section “*Analysis: synaptic sign index*”). Synaptic sign indices of -1, 0, and 1 indicate pure excitation, balanced input (i.e., the amplitude of the EPSC and IPSC are equal), and pure inhibition, respectively. In total, 28 cells (from 10 mice) were recorded from animals at baseline, 88 cells (from 21 mice) were recorded from animals after reward pairing, and 61 cells (from 14 mice) were recorded from animals after punishment pairing. Cells in all three groups had similar passive properties and there are no significant correlations between synaptic sign index and recording parameters (**fig. S10**). Using data from all recorded cells, no clear differences emerge in synaptic sign, as reflected in the synaptic sign index, across baseline, reward pairing, and punishment pairing groups (**fig. S11, S12A, B**).

**Fig 3.**
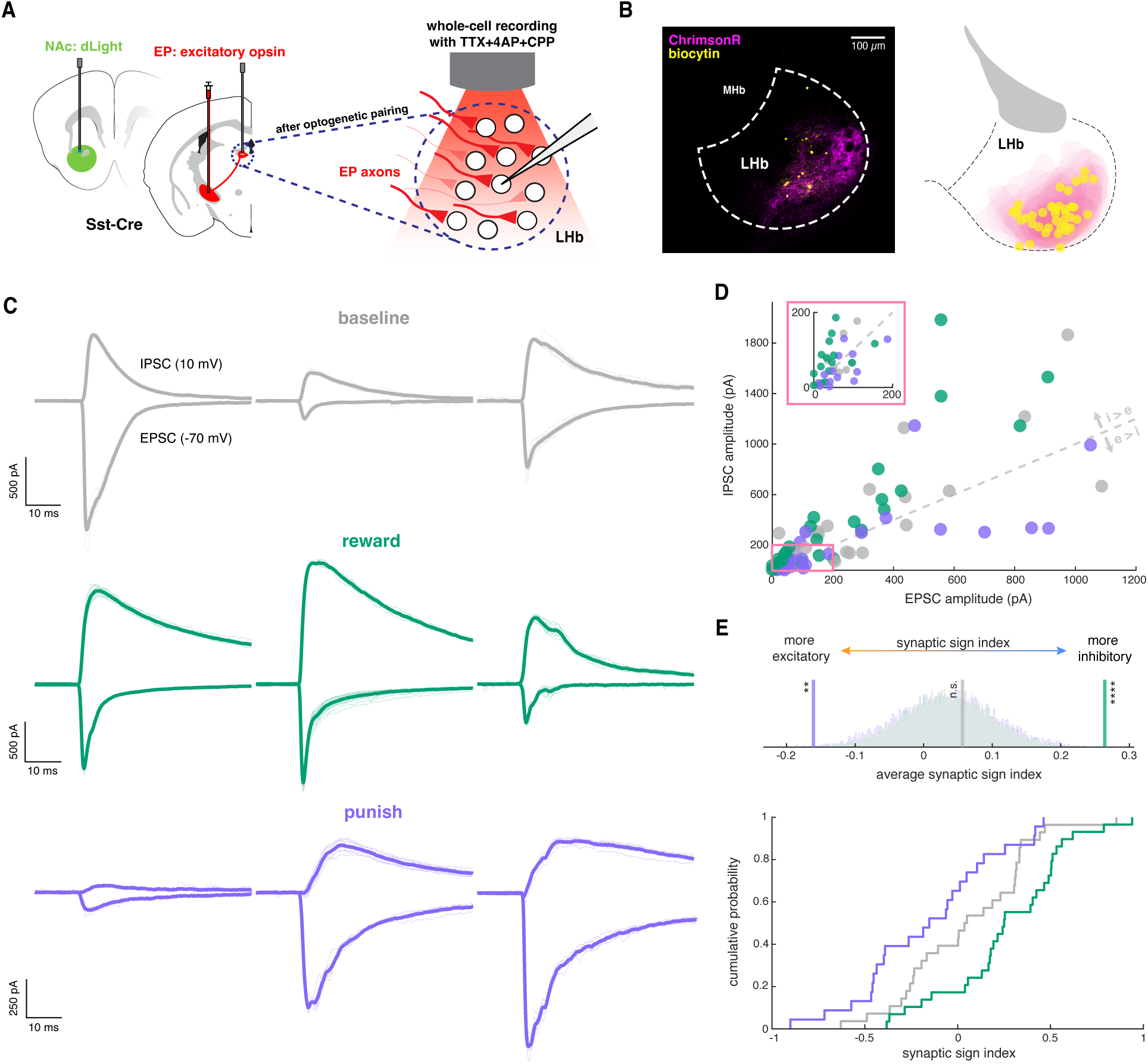
Sign switching occurs at EP Sst**→**LHb synapses during opto-outcome reversal learning **A**, Overview of experiment. We selectively express excitatory opsin in EP Sst neurons (see Methods, n=44 mice). Animals may undergo reward or punishment pairing while NAc DA transients are recorded (34/44 mice). Acute brain slices of LHb are then obtained to perform whole-cell voltage-clamp recordings on LHb neurons. Monosynaptic inputs from EP Sst neurons to LHb neurons are measured by optogentically stimulating opsin-expressing EP axons in LHb. **B**, Example histology (*left*) and locations of biocytin expressing cells with red shading indicates approximate spread of EP Sst axons in LHb of the recovered slices (*right*). **C**, Example cells for animals undergo baseline (gray), punish-to-reward pairing (green), and reward-to-punish pairing (purple) condition. Thicker lines indicate the averaged PSCs from all excitatory or inhibitory sweeps of this cell and thinner lines indicate individual sweeps. **D**, Scatter plot of EPSC amplitude vs IPSC amplitude (baseline: n=28 cells, 10 mice; punish-to-reward: n=29 cells, 7 mice; reward-to-punish: n=23 cells, 5 mice). Each dot represents a cell. Gray dotted line indicates where EPSC amplitude is equal to IPSC amplitude. Pink inset shows a zoom-in view of cells with smaller EP Sst neuron inputs. **E**, Synaptic sign index of the three conditions. *Top*, synaptic sign index is a cell-based metric that measures the difference between inhibitory and excitatory current amplitude normalized by total amount of input currents for each cell. Synaptic sign index closer to 1 indicates an inhibitory biased EP Sst→LHb synapses received by the recorded LHb neurons and sign index closer to -1 indicates an excitatory biased EP Sst→LHb synapses. *Middle*, average synaptic sign index of all three conditions (vertical lines) and the corresponding bootstrapped distributions used to calculate significance (semi-transparent bars). *Bottom*, cumulative distribution function of synaptic sign index of each condition.

However, when we restricted the analysis to data from cells recorded on days on which the opto-outcome pairing was reversed (i.e., switched from punishment to reward or vice versa; see Methods and **fig. S9**), differences in synaptic sign index across groups became evident. On reversal days, postsynaptic currents in LHb cells evoked by EP Sst axons shifted toward more inhibitory transmission after transition from punishment to reward (n=29 cells, 7 mice), and toward more excitatory transmission with transition from reward to punishment (n=23 cells, 5 mice) (**Fig. 3C-E**; **fig. S12C, D**). Moreover, presentation of water or air puff alone to thirsty animals did not induce sign switching between EP Sst→LHb synapses, indicating that changes in synaptic sign index are not driven by the outcome per se but instead require updating of experimentally imposed valence on EP Sst neuron activity (**fig. S13**).

Thus, synaptic sign index reveals changes in excitatory vs inhibitory synaptic transmission in a population of EP Sst→LHb synapses induced by *in vivo* opto-outcome reversal learning, consistent with reversal of the valence associated with the EP opto stim triggering synaptic sign switching at these synapses.

## Synaptic sign of EP Sst**→**LHb synapses correlates with recent dopamine updates

Since we observed pairing-induced changes in synaptic sign index on the day of opto-outcome reversal but not across all sessions, we hypothesized that variability in learning within individual sessions, as reflected in experience-dependent changes in DA signaling, may explain this discrepancy. DA dynamics fluctuate on a trial-by-trial basis, potentially due to uncontrolled sensory stimuli or changes in internal state such as stress or thirst satiation, which in turn may affect the perceived value of stimuli.

To examine potential links between EP Sst→LHb synaptic sign and DA, we calculated summary metrics for the *ex vivo* synaptic sign index and the changes in *in vivo* EP opto stim-evoked DA transients across trials for each animal (**Fig. 4A**). To summarize in-session changes in DA (**Fig. 4A**, *left*), we measured the slope of changes in the amplitude of EP opto stim-evoked DA transients across trials in the last behavioral session. We calculated the slope of DA amplitude vs. trials at the beginning (first 20 trials) and end (last 20 trials) of the session (referred later as early or late DA slope). To summarize the synaptic sign index per animal (**Fig. 4A**, *right*), we averaged the synaptic sign indices from all recorded cells of each animal, which we referred to as animal sign index. Since LHb di-synaptically inhibits VTA and NAc DA release, animals with increasing DA responses over trials might be expected to have inhibitory-biased EP-LHb synapses (higher animal sign index) whereas those with decreasing NAc DA responses might be expected to have excitatory-biased synapses (lower animal sign index). Thus, we predict a positive correlation between animal sign index and DA slope (**Fig. 4B**).

**Fig 4.**
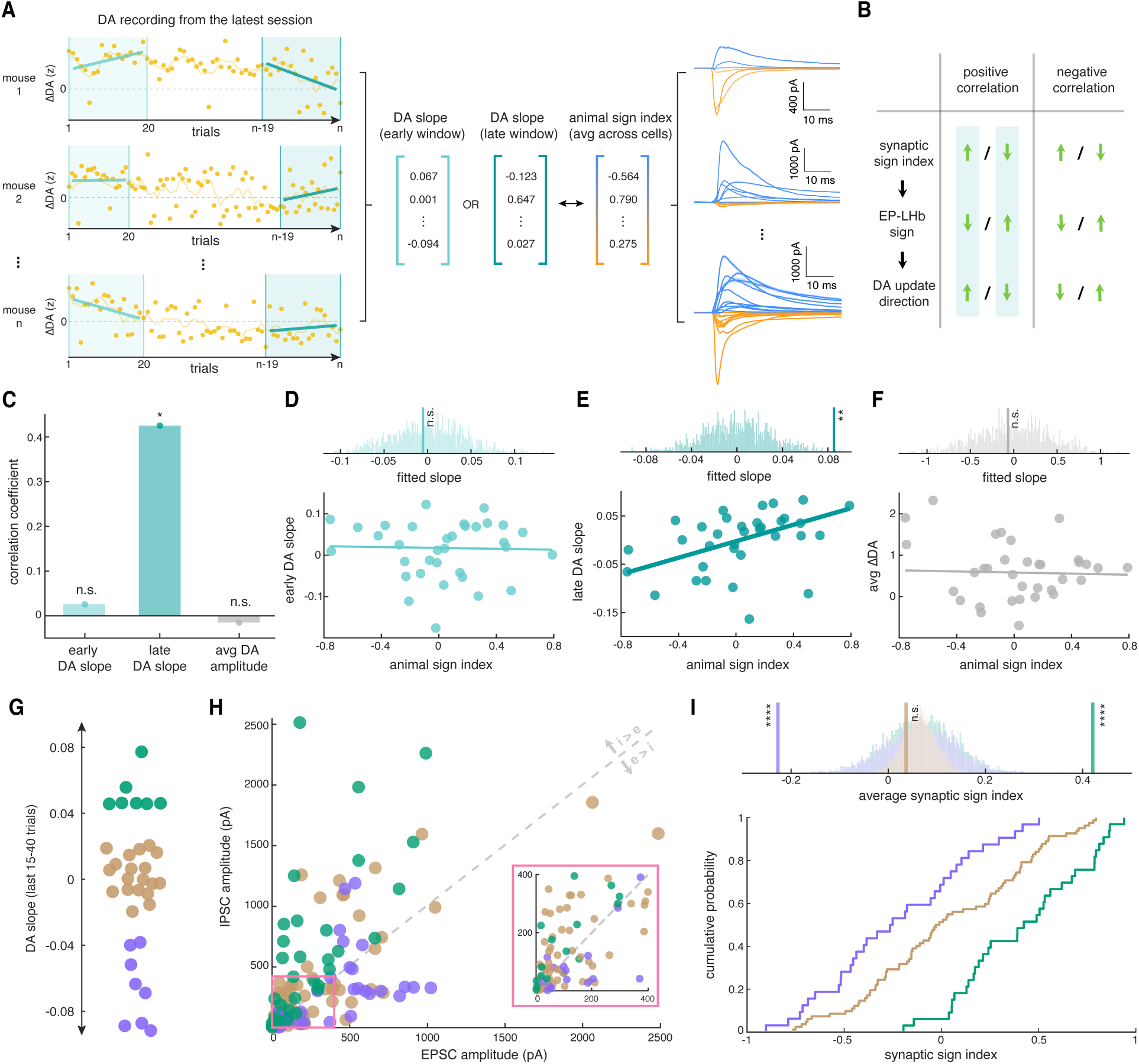
Synaptic sign of EP-LHb strongly correlates with recent dopamine updates **A**, Overview of analysis. For each animal (n=34 mice), we extracted DA amplitudes in response to EP opto stim in all opto-only trials of the last recording session. We define an early and late window within the session that correspond to the first 20 and the last 20 trials. We then calculated the slope of DA amplitude vs trials (smoothed using the nearby 5 trials) for both early and late window. This generates two vectors for early and late window summarizing the change of evoked DA amplitude of each animal during these two windows (termed “DA slope”). On the other end, we also calculate the “animal sign index” vector by averaging synaptic sign index of all cells recorded from that animal. We analyze the relationship between the animal sign index vector and DA slope vectors. **B**, Summary of possible direction of correlation between animal sign index and DA slope correlation and corresponding interpretations. Based on anatomy of the EP-LHb-VTA circuit, described in **Fig. 1A**, a positive correlation would be expected (light blue shade). **C**, Correlation coefficients calculated between animal sign index and either early DA slope (light green), late DA slope (dark green), or session-averaged DA amplitude to opto-only trials (gray). Correlation coefficients are calculated using Spearman’s rank correlation. **D**, Correlation between animal sign index and early DA slope. *Top*, fitted slope (vertical line) and distribution build from bootstrapped simulations of the fitted slope used to calculate significance (semi-transparent bars). *Bottom*, scatter plot of animal sign index vs early DA slope. Each dot represents each animal. Line indicates the fitted linear function used to calculate the slope. **E**, Correlation between animal sign index and late DA slope. **F**, Correlation between animal sign index and averaged DA amplitude. **G**, Classification of fitted DA slopes during late window. Each dot represents an animal. K-means clustering was repeated 100 times to separate animals into 3 groups indicating increasing, stable, and decreasing DA amplitude during the late window (average of last 15 to last 40 trials). **H**, Scatter plot of EPSC amplitude and IPSC amplitude of all cells recorded. Each dot represents a cell. Cells are colored based on whether the animal’s evoked DA amplitude is increasing (green, n=33 cells, 7 mice), stable (brown, n=82 cells, 19 mice), or decreasing (purple, n=32 cells, 8 mice) during the late window. Pink inset shows a zoom-in view of cells with smaller EP Sst neuron inputs. **I**, Synaptic sign index of the three conditions of increasing, stable, or decreasing DA amplitude. *Top*, average synaptic sign index of all three conditions (vertical lines) and the corresponding bootstrapped distributions used to calculate significance (semi-transparent bars). *Bottom*, cumulative distribution function of synaptic sign index of each condition.

We first computed the spearman correlation coefficients of animal sign index with the early and late DA slopes as well as with the average amplitude of EP opto stim-evoked DA transients.

Only the slope during the late window correlates significantly with animal sign index (**Fig. 4C-F**). To reveal stretches of trials in which changes in DA amplitude *in vivo* correlates most strongly with animal sign index measure *ex vivo*, we varied the length of early and late windows (**fig. S14A**). We find that the animal sign index significantly correlated with the slope of DA amplitude changes in last 15 to 40 trials (**fig. S14B**), corresponding to approximately 6–20 minutes before the end of the session (**fig. S14C**). The slope of DA amplitude changes during the early window and average DA amplitude in the analysis windows were not consistently significantly correlated with per animal sign index (**fig. S14B**). To test for spurious correlations specific to this dataset, we randomly separated our data into training (27 mice, i.e. 80%) and test (7 mice, i.e. 20%) sets and shuffled the groups 200 times (**fig. S15A**). For each shuffle, we identified stretches of trials during which the slope of changes in DA amplitude significantly correlates with sign index for animals in the training set and then tested the strength of the correlations in the test set (**fig. S15A**). Across all 200 randomly selected train-test splits, spearman correlation coefficients between the slope of changes in DA amplitude during late window and sign index is significantly higher than those calculated from DA amplitude changes in early window or average DA amplitudes (**fig. S15B**). Together, these findings demonstrate that sign of EP-LHb synapses is tightly linked to the direction of DA updates in the final ∼20 minutes of behavior (**fig. S14C**), supporting the notion that EP Sst→LHb synaptic plasticity correlates with recent learning-associated DA updates.

Since EP-LHb synaptic sign correlates with changes in the EP opto stim-evoked DA transients near the end of the behavioral session, we examined if grouping data from all animals and sessions based purely on changes in the slope of DA transients near the end of the session would reveal differences in synaptic sign index across groups. This analysis ignores the direction of the EP opto stim-outcome pairing, but instead tests only if recent changes in DA amplitude correlate with sign index. For each animal, we calculated the average late DA slope across the last 15 to 40 trials in the last session before *ex vivo* analysis (**fig. S14B**). We used K-means clustering to divide late DA slopes from all animals into one of three groups, corresponding to animals with increasing (green), stable (brown), and decreasing (purple) amplitude of opto-evoked DA transients during the late window (**Fig 4G**). Separating all recorded cells based on this per animal *in vivo* criterion reveals differences in synaptic sign across groups that grouping by reward/punishment pairing (**fig. S11**) did not (increasing DA: green, n=33 cells, 7 mice; stable DA: brown, n=82 cells, 19 mice; decreasing DA: purple, n=32 cells, 8 mice; **Fig 4H, I**; **fig. S16**). These correlations are maintained in randomly selected test sets not used for training (**fig. S15C, D**) and are not solely driven by data collected on days of opto-outcome reversal, as sign switching persists even when removing these cells from analysis (**fig. S17**).

Together, these results demonstrate that the sign of EP Sst→LHb synapses shifts in tandem with changes in the valence of the context in which their activity is paired: synapses become more inhibitory when DA release is increasing and more excitatory when DA release is decreasing over time. The correlation between EP Sst→LHb synaptic sign and DA updates is only seen within a short time window (∼20 minutes), suggesting that sign switching at these synapses is a unique experience-dependent plasticity that operates at fast behavior timescales to mediate online DA updates and support ongoing learning.

## Discussion

Our study reveals a previously undescribed form of synaptic plasticity in the mammalian brain: rapid experience-dependent synaptic sign switching. This plasticity occurs at the output of EP Sst neurons, which co-package and co-release glutamate and GABA, and is expressed by changes in the relative contributions of glutamatergic and GABAergic currents evoked in their LHb targets. Furthermore, *in vivo* pairing activity of EP Sst→LHb synapses with updates in value, bi-directionally modulate behavior and the effects of EP Sst activity on DA release in NAc, supporting the physiological importance of these mechanisms. Our results reveal that EP Sst→LHb synaptic plasticity, which includes synaptic sign switching, is the circuit-level mechanism for updating the relationship between EP Sst activity and NAc DA in an experience-dependent manner.

The finding that synapses can quickly switch signs challenges a long-held assumption of mammalian brains. Traditional models of activity-dependent synaptic plasticity posit that learning modulates synaptic strengths but not synaptic signs. Previous studies have described neurons or axonal branches whose neurotransmitter identity switches on long time scales (days to weeks) (*35–37*). Furthermore, changes in mRNA expression in these examples seem to underlie a cell-wide switch in neurotransmitter identity (*36, 37*). However, synaptic sign-switching between excitation and inhibition within the timescales of minutes has not been previously demonstrated. Moreover, neurons that co-express Vglut2 and Vgat (*18*) and biphasic glutamatergic and GABAergic synaptic currents (*38*) suggestive of glutamate and GABA co-packaging are found in many brain regions, such that the phenomena revealed here may not be rare and may occur at synapses outside of the LHb.

Studying the relationship between synaptic plasticity and behavior is challenging because it is difficult to isolate synaptic currents *ex vivo* that underwent plasticity *in vivo*. Furthermore, synaptic efficacy is difficult to repeatedly measure *in vivo* during learning. By pairing the optogenetic activation of EP Sst neurons with positive and negative outcomes, we can assign a valence to the activity of these glutamate and GABA co-releasing cells and thereby address both challenges. First, by using optogenetic activation of the same synapses *in vivo* and *ex vivo*, we can reliably induce synaptic plasticity *in vivo* and optogenetically re-activate these projections in acute brain slices to measure synaptic signs *ex vivo*, allowing us to both induce plasticity and measure synaptic states in a cell-type and synapse-specific manner. Second, because the LHb has a fixed inhibitory effect on NAc DA release, we were able use the modulation of optogenetically evoked release of DA in NAc as a surrogate for the experience-dependent modulation of EP Sst→LHb synapses *in vivo.* Using this strategy, we demonstrate the existence of experience-dependent plasticity that shifts the net balance of excitation and inhibition at genetically-defined and physiologically-relevant synapses, producing functional switch in synaptic signs.

Mechanistically, we provide evidence that postsynaptic signaling is crucial for EP Sst→LHb synaptic sign switching: the effects of opto-outcome pairing on NAc DA release and behavior are abolished when LHb neurons are lesioned or when postsynaptic CaMKII activity is inhibited. Furthermore, glutamate release induced by optogenetic stimulation of EP co-releasing neurons remains unchanged even as the relationships between the optogenetic stimulation and both DA release and behavior change, consistent with results observed by others (*31*). These results disfavor the possibility that sign switching at EP Sst→LHb synapses occurs via purely presynaptic mechanisms, as this would require large increases or decreases in GABA content per vesicle without affecting glutamate levels—an unlikely scenario given vesicle volume constraints and the competitive equilibrium might govern the neurotransmitter loading in these synapses (*39, 40*). On the other hand, postsynaptic plasticity of glutamate and GABA receptors in LHb neurons has been shown to mediate punishment- and reward-related learning, respectively (*27, 29, 31*). Therefore, experience-dependent sign switching at EP Sst→LHb synapses may be driven primarily by postsynaptic mechanisms, such as differential activity-dependent insertion of glutamate and GABA receptors.

Our results offer insight into the potential timescale with which plasticity is induced and behaviorally relevant: the sign of EP Sst→LHb synapses significantly correlates with DA updates that occurred 6-20 minutes before the end of the session but not with longer-term behavioral or experimental variables. Levels of cue-evoked DA release often fluctuate within a single session, possibly driven by changes in external environment or internal bodily states (e.g. value of the reward-predicting cue devalues as the animal becomes satiated and thirst diminishes). Categorizing sessions purely based on the direction of opto-outcome contingency ignores these crucial fluctuations that occur on a shorter timescale. This is consistent with our initial observation that EP Sst→LHb synaptic signs do not differ when observations are grouped by the direction of opto-outcome pairing. However, categorizing sessions based on direction of NAc DA updates on a shorter timescale (i.e., the most recent ∼20 minutes or ∼40 trials) reveals the relationship between synaptic sign and changes in NAc DA: EP Sst→LHb synapses are more inhibitory when cue-evoked DA transients are increasing and more excitatory when DA transients are decreasing during this window.

This correlation between EP Sst→LHb synaptic sign and DA updates supports a model in which these sign-switching synapses react to feedback, that likely resembles RPE, to tune DA transients, offering a rapid update signal to allow the brain to track changes in the environment. The nature and source of this hypothesized feedback error signal is currently unknown.

Moreover, because of delays in preparing acute brain slices and the need to aggregate data across multiple trials (> 10 trials) to generate a confident estimation of DA updates, we may have underestimated the speed of sign-switching plasticity. In addition, blocking CaMKII in LHb prevents learning dependent changes in NAc DA signals, suggesting involvement of classical plasticity inducing pathways (*32, 41*).

Despite our demonstration of the existence of synaptic sign switching in LHb, the segregation of excitatory and inhibitory transmission and the fixed sign of synaptic output likely holds in most of the brain. It is interesting to speculate why other synapses in the brain do not implement such plasticity and why sign switching is beneficial at EP Sst→LHb synapses. In ANNs, the strength of each synapse is typically set using gradient descent to minimize a cost function, which is roughly equivalent to minimizing the error in the calculation carried out by the circuit. Gradient descent in multi-layer networks, such as the brain, requires that each synapse be updated based on a rule that requires knowing how the activity of both the pre- and post-synaptic neuron relates to the error, which is difficult to implement in biological circuits (*1*). To circumvent this limitation, synapses in the brain may be constructed along positive and negative channels, each with fixed relationship to a locally calculated error (*42*).

On the other hand, the computational responsibilities of the LHb and its fixed inhibitory relationship with NAc DA neurons may both necessitate and permit fast sign-switching plasticity that implements gradient descent. The EP Sst→LHb→NAc DA circuit operates essentially as a simple three-layer network in which the plasticity of inputs (EP) onto a hidden layer (LHb) control the output (NAc DA). In such an architecture in which the sign between the hidden layer and the output is fixed (negative in this case), the local activity of each EP–LHb neuron pair can be directly tied to the overall error, making a global error signal sufficient to implement gradient descent.

Animals adapt to changing environments through flexible mapping of actions to outcomes. Yet, this mapping is context dependent and plastic such that the same action may be predictive of a positive outcome in one context but negative outcome in the other, requiring animals to dynamically estimate the value of the expected outcome of carrying out an action. The basal ganglia funnels information from ∼17 million corticostriatal neurons into ∼1000 EP neurons, representing an enormous convergence of cortical signals (*43–45*). Thus, the activity of inputs to EP is enriched with information about external and internal state that is likely relevant to action selection (*7, 8*). EP Sst neurons relay this information to LHb for calculation of expected value and modulation of VTA DA neurons that signal RPE. Therefore, the sign-switching property of EP Sst→LHb synapses and their update rules revealed here provide a natural mechanism that ensures value updates can be rapidly and faithfully calculated with no potential computational ceiling (*3*).

## Acknowledgments

We thank the Sabatini laboratory and M. Wallace, M. Andermann, F. Wang, C. Harvey, J. Assad, and N. Uchida for helpful discussions. We thank the Core for Imaging Technology & Education for imaging help. We thank T. Pottala, A. Biwas, Y. Dong, C. King, M. Wen for mouse husbandry and genotyping. We thank E. Drewry, C. Walker for laboratory administration.

## Author Contributions

SL and BS designed the experiments. SL processed data. SL conducted the analyses, designed the behavior rig, and performed surgeries. SL, GK, EJ, CZ and EB trained mice and performed fiber photometry recordings. WW performed *ex vivo* electrophysiology. PC cloned and characterized the WiChR virus and helped with *ex vivo* electrophysiology. SL and BS wrote the manuscript.

## Data and materials availability

All data needed to evaluate the conclusions in the paper are present in the paper and/or supplemental materials.

**Supplementary Materials** Materials and Methods Figs. S1 to S1

## Materials and Methods

All procedures were carried out in accordance with the IACUC protocol #IS00000571-6. All animal care and experimental manipulations were performed in accordance with protocols approved by the Harvard Standing Committee on Animal Care, following guidelines described in the US NIH Guide for the Care and Use of Laboratory Animals.

### Mice

The following mouse lines were used: C57BL6/J (The Jackson Laboratory, 000664); *Sst-IRES-Cre* (The Jackson Laboratory, #013044); *Vglut2-IRES-Cre* (The Jackson Laboratory, #016963); and *Vgat-IRES-Flp* (The Jackson Laboratory, #029591). All mice were bred on a C57BL/6J genetic background and heterozygotes were used unless noted. For behavioral experiments, 6 to 15-week-old mice were used. We used male and female mice in an approximately equal ratio for all experiments. We did not observe any differences in learning between different sexes. All figures include data from both males and females.

### Viruses

The following are all viruses used, with source and titer during injection indicated in parentheses:

- AAV8-hSyn-FLEX-ChrimsonR-tdTomato (UNC Vector Core, 3.9E12 GC/ml)
- AAV9-hSyn-dLight3.8 (Boston’s Children’s hospital core (BCH); 1E13 GC/ml)
- AAV9-hSyn-dLight1.1 (Boston’s Children’s hospital core (BCH); 1E13 GC/ml)
- AAV8-nEF-Con/Fon-hChR2(H134R)-EYFP (Addgene, 1E13 GC/ml)
- AAV8-hSyn-ConVERGD 2.0-ChRmine-oScarlet (Custom production, 1E13 GC/ml)
- AAV1-hSyn-SF-iGluSnFR.A184S (Addgene, 1E13 GC/ml)
- AAV5-Ef1a-FLEX-taCasp3-TEVp (Addgene, 1E13 GC/ml)
- AAV8-CaMKIIP-mEGFP-P2A-paAIP2 (Addgene, 1.2E13 GC/ml)
- AAV9-CAG-FLEX-WiChR-oScarlet-WPRE (Custom production, 1.05E12 GC/ml)
- AAV9-CaMKII(0.4kb)-Cre-SV40 (Addgene, 1.8E13 GC/ml)

To stimulate the glutamate and GABA co-releasing neurons in EP, we mainly expressed Cre-dependent opsin ChrimsonR in *Sst-Cre* animals. Since Sst neurons are the only population in EP that co-expresses Vglut2 and Vgat, in a subset of animals in Fig. 3, 4 (13/44 mice), Sst neurons were targeted by injecting viral constructs that drives ChR2 or ChRmine expression under the presence of both Cre and Flp in *Vglut2-Cre x Vgat-Flp* animals. To achieve specific expression for ChRmine only under presence of both Cre and Flp recombinase, we cloned the ChRmine-oScarlet gene into the ConVERGD construct (*46*). All recombinase-dependent viruses were injected in wild-type animals at the region of interest to confirm the lack of leaky (i.e., recombinase independent) expression (**fig. S2**).

### Intracranial injections: coordinates

The following coordinates were used to deliver viruses to region of interest. The volume of injection is indicated in parentheses:

- EP: -1.10 mm A/P, ±1.95 mm M/L, -4.45 mm D/V (150 nl)
- LHb: -1.65 mm A/P, ±0.46 mm M/L, -2.85 mm D/V (150 nl for FLEX-taCasp3, 200 nl for other viruses)
- NAc: 1.55 mm A/P, ±1.40 mm M/L, -4.25 mm D/V (300 nl)
- TRN: -1.10 mm A/P, ±1.95 mm M/L, -3.80 mm D/V (50 nl)

### Intracranial injections: surgery procedure

Mice were anaesthetized with 5% isoflurane and maintained during surgery with 1.5% isoflurane and 0.08% O_2_. Under the stereotaxic frame (David Kopf Instruments), the skull was exposed in aseptic conditions, a small craniotomy (around 300 μm) was drilled, and virus was injected into the targeted regions. Injections were done by first lowering a pulled pipette to 100 µm deeper than the targeted region, retracted to the targeted coordinate and waiting for 3 min. Using a syringe pump (Harvard Apparatus, 883015), virus was infused at the rate of 30 nl/min for EP and LHb, and 50 nl/min for NAc. After the infusion was completed, the pipette remained in place for a further 5 min, then retracted slowly with at least 3 cycles of slow withdrawal of 50 µm at 1 min intervals. All virus injections were performed unilaterally.

For fiber implants, after AAV injection, the skull was scored lightly with a razor blade to promote glue adhesion. Then, a 200-µm blunt-ended fiber (Doric Lenses, MFC_200/230- 0.48_4.4 mm for NAc and MFC_200/230-0.48_3 mm for LHb) was slowly inserted into the brain until it reached 100 µm above the injection site. For iGluSnFR recording in LHb, a 4.4 mm long 200 um blunt-ended fiber (Doric Lenses, MFC_200/230-0.48_4.4 mm) was implanted from the opposite side of the brain with a 45-degree angle. The implantation coordinate is ±1.95 mm M/L, -2.65 mm A/P, and 3.53 mm deep from the brain surface. The fiber was held in place with glue (Loctite gel, 454) and hardening was accelerated with the application of Zip Kicker (Pacer Technology). A metal headplate was glued at lambda and white cement (Parkell) was applied on top of the glue to further secure the headplate and fibers.

After the surgery, mice were placed on heating pad until their activity recovered and then returned to their home cage. Mice were given pre- and post-operative oral carprofen (CPF, 5 mg per kg per day) as an analgesic and monitored daily for at least four days after surgery. At least 17 days passed after virus injection before experiments were performed.

### Behavior: behavior setup

Custom written code in Arduino (MEGA 2560) was used to deliver stimuli and detect animal behavior. Critically, the Arduino also delivers an irregular sync pulse that is used to synchronize with the recording systems (Labjack for photometry, SpikeGLX for behavioral data). The sync pulse consisted of a 50 ms square pulse delivered at a random 100-500 ms intervals throughout the recording session.

Licks were detected by recording the voltage drop between the spout and the tube. The inside of the tube was covered with copper foil and grounded. Solenoids (The Lee Company, LHQA0531220H) were connected to 20-ml syringes, acting as water reservoirs, and opened for a short duration to deliver water rewards. Water reward size was calibrated by adjusting solenoids opening time (∼20 ms). The water delivery spout was made by connecting blunt syringes needles (18 gauges) to the solenoids via tubing (Cole-Parmer, EW-06460-34). A speaker (Madiasound, tw025a20) connected to an amplifier (FOSTEX, AP05) was positioned underneath the tube and connected to the Arduino to deliver tones during the task. All behavioral events and stimuli were copied to and acquired in SpikeGLX along with neural recording data. A 473 nm or 637 nm laser was used to deliver blue or red stimulation, respectively.

### Behavior: fiber photometry setup

Fiber implants on the mice were connected to a 0.48 NA patch cord (Doric Lenses, MFP_200/220/900-0.48_2m_FCM-MF1.25, low autofluorescence epoxy), which received excitation light and propagated its emission light. For NAc dLight recordings, a 470 nm LED (Thorlabs, M470F3) and a LED driver (Thorlabs, LEDD1B) delivered blue excitation light and a Doric filter cube (Doric Lenses, FMC5_E1(465–480)_F1(500-540) _E2(555-570)_F2(580- 680)_S) filtered and detected 500-540 nm green emission light. For LHb recordings, a Doric filter cube was used to both deliver excitation light and detect emission light (465–480 nm blue excitation light; 500–540 nm green emission light). Excitation light was amplitude-modulated at 200 Hz and 250 Hz using MATLAB. Signals from the photodetectors were amplified in DC mode with Doric amplifiers and received by a Labjack (T7) streaming at 2,000 Hz. The Labjack also receives the Arduino generated sync pulse.

### Behavior: water restriction and habituation

Singly housed mice were restricted to 1 ml water per day before training and were maintained at > 80% of their initial body weight for the full duration of training and photometry. All training sessions were conducted under red light.

We habituate mice to the head-fixed setup using the following schedule. In days 1-3, we habituate animals to a gloved hand in the home cage and gradually moved them outside the home cage for 15 min while delivering water. At day 4-5, when animal is comfortable to being handled, we begin head-restraint.

At initial stage of head-restraining, we manually deliver water through the lick spout, which is positioned around 1 mm to the mouth. Once the mouse starts to lick, we manually deliver several more water droplets during each licking bout to encourage licking behavior. After spontaneous licking to the spout is observed, we switched to randomly deliver the reward at intervals defined by a randomly selected enforced non-lick (ENL) window (see below for more detail). Animals were taken out of the head-fix setup upon satiation (i.e. they stop licking even when water was delivered). After each session, water is supplemented such that the total amount of water is ∼1 ml for each day. Optical fibers are attached during habituation to mimic the recording conditions.

### Behavior: general recording schedule

Recording session are divided into three main phases. A baseline phase in which different stimuli are given at random with no contingency, and reward or punish phases in which optogenetic stimulus were paired with reward (water) or punishment (air puff), respectively. Each session ends when the animal no longer obtains free water or reaches ∼50 minutes on the setup. Reward and punishment pairing phases normally last for three days, but the exact duration depends on the speed of learning for each animal. Recording always starts with baseline sessions, then reward sessions. After reward pairing sessions, animals might progress to punish pairing sessions or be sacrifice for *ex vivo* electrophysiology (**fig. S9**). Details of each session will be elaborated below. Before each recording, laser power exiting the optical fiber is calibrated to a target value using a power meter (Thorlabs, PM100D).

### Behavior: enforced non-lick window (ENL)

For all three phases of recording, an ENL window is used to decide when to deliver the next stimuli. Animals must withhold licking during the ENL window, which is randomly chosen between 2-4 seconds. If animals lick during ENL window, the time will be reset until it succeeded to withhold licking. In addition, time interval between trials are always more than 15 seconds to ensure recovery of neural signal and behaviors to baseline. This means that, if the animal withhold licking for 2-4 seconds but it has been less than 15 seconds since the start of the previous trial, the animal must wait until 15 seconds have passed while continues to withhold licking to enter the next trial.

### Behavior: baseline sessions

Once animals show robust licking behavior to the spout, the transition to the “baseline” recording phase. In this phase, four stimuli are delivered at random with no contingency between them. The following are delivered with the indicated probabilities:

- 40%: 2 µl water reward,
- 10%: 200 ms air puff
- 40%: 500 ms optogenetic stimulation targeted to EP (or TRN in the case of **fig. S5**)
- 10%: 500 ms continuous tone (3000 Hz) that is ∼20 dB above the environmental noise (∼65 dB)

Optogenetic stimulations of EP using ChrimsonR or ChRmine consist of 5 ms, 8 mW pulses delivered at 50 Hz. For Con/Fon-ChR2, optogenetic stimulation of EP consist of 5 ms pulse delivered at 30 Hz at 3 mW. After 3-4 days of baseline session, animals were progressed to reward pairing session and then punishment pairing sessions in which the optogenetic stimulus was paired to a water reward or an air puff punishment (see below).

### Behavior: reward or punishment pairing sessions

In both reward and punishment pairing sessions, all trials start with a 500 ms cue period during which one of two types of cues will be presented in most trials and paired with outcome in a deterministic fashion. In roughly 50% of trials, the opto-only cue will be delivered. In roughly 40% of trials, the opto+tone cue will be delivered. The opto+tone cue consists of the standard optogenetic stimulation plus the 500 ms continuous tone. In the remaining trials, a tone-only cue will be delivered which consists of only the continuous tone without concurrent optogenetic stimulation. Tone-only cues always lead to outcome omission.

In reward pairing sessions, both opto-only and opto+tone cues are deterministically associated with a water reward. However, the amount of reward will be different based on animal’s action during the response period (500 ms to 2000 ms). If the animal performs more than 2 licks during the cue and response period, a large 10 µl water reward is delivered. Otherwise, a small 2 µl water reward is delivered to the animal at the onset of free outcome period (2000 ms after the current trial start). If the animal successfully obtained the large reward, the inter-trial interval (ITI) will be a normal ENL window (see above). However, if the animal fails to get the large reward, a 10 s timeout period is added on top of the normal ENL window, resulting in a long ITI.

In punishment pairing sessions, both opto-only and opto+tone cues will be deterministically associated with an air puff. As in the reward pairing sessions, the duration of air puff is different based on animal’s action during the response period (500 ms to 2000 ms). If the animal performs more than 2 licks during cue and response period, a large 200 ms air puff is delivered. Otherwise, a small 100 ms air puff is delivered to the animal at the onset of free outcome period (2000 ms to next trial start). In 10% of trials, an un-cued 2 µl water reward is delivered to the animal to maintain engagement to the task. In some mice (e.g. mice in Fig. 1), a short air puff is delivered unconditionally at 120 ms after cue offset. If the animal obtained the short air puff, the inter-trial interval (ITI) will be a normal ENL window (see above). However, if the animal gets a large air puff, a 10 s timeout period is added on top of the normal ENL window, resulting in a long ITI.

### Behavior: preventing mice from cheating

To prevent the possibility that the mouse using opto-related sensory stimuli as a cue rather than the optogenetic activation of the neuron population, several features are implemented in the behavior setup. First, we placed an overhanging red lamp around 10 cm above the mouse head to mask the laser light leakage through the connection between the fiber and patch cord (**movie S1**). Second, we use a silent galvanometer scanning mirror to direct the light in or out of a pin hole to deliver optogenetic stimulation, such that no sound would be generated during optogenetic stimulation. Lastly, no opsin expression controls were performed in which we deliver light to the fiber and LHb but the tissue does not express an opsin (**fig. S4**).

### Behavior: taCasp3 ablation of LHb neurons

Animals in the taCasp3 ablation cohort were trained normally. No motor and gross behavior anomalies were observed. Animals first went through baseline, reward, and punishment pairing sessions as usual. Afterwards, they underwent reward then punishment pairing sessions in which the auditory tone was presented in > 80% of trials.

### Behavior: paAIP2 inhibition of CaMKII in LHb

Animals in the taCasp3 ablation cohort were trained normally. No motor and gross behavior anomalies were observed. Animals first went through baseline, reward, and punishment pairing sessions as usual. For all reward and punish pairing sessions, 1 s continuous light pulse at 10 mW was delivered at 5 s intervals throughout the session to enable constant inhibition of CaMKII. Animals then underwent opto-reward pairing sessions without activation of paAIP2.

### Behavior: WiChR inhibition of LHb

For WiChR-mediated inhibition of LHb, we deliver continuous light pulse for 500 ms at ∼ 0.3 mW using a 473 nm laser that delivers light through the same fiber on LHb. For recording sessions of opto pairing, 15 blue light pulses (500 ms, continuous) were delivered to the animal after stopping the task. Triggers for the blue light pulses are produced by an Arduino. We tried our best to avoid delivering light pulses at the time of any spontaneous dopamine increase, as doing this will affect the amplitude of WiChR-induced dopamine transients, but due to their unpredictability, ∼50% of trials occurred overlapped with spontaneous dopamine transients.

### Behavior: from in vivo opto-pairing to ex vivo electrophysiology

To comprehensively assess how EP Sst→LHb synaptic sign might relate to dopamine updates, different animals were transferred for *ex vivo* electrophysiological analysis at different phases of the task. In general, all animals experienced baseline and reward pairing sessions to ensure that they acquired the opto-outcome association. Afterwards three types of schedules were used, as illustrated in below and in **fig. S9**.

Before electrophysiological analysis, all mice experienced a complete punishment or reward pairing phase (∼3 sessions) before the opto-outcome contingency was reversed. Subsequently, acute brain slices were prepared either on the day of the reversal (day 1 data), 2-3 days after reversal (day 2-3 data), or more than 3 days after reversal (3+ day data). Behavioral sessions on the day of acute slice preparation contained at least 20 opto-only trials. After finishing *in vivo* opto-pairing, animals were put back into their home cage, covered with black plastic bags and placed in a dark room until processing. In addition, most acute brain slices were cut less than 30 min after the end of recording session.

For control experiments that involves delivering only reward or punishment outcomes with no EP opto stim pairing (**fig. S13**), animals were water deprived, handled and habituated the same way as other animals illustrated above. However, they were only exposed to reward outcome (water) or punishment outcome (air puffs) on the day of *ex vivo* recordings.

### Ex vivo electrophysiology: preparation of acute brain slices

Acute brain slices were prepared from adult mice anesthetized by isoflurane inhalation and perfused transcardially with ice-cold, carbogen-saturated artificial cerebral spinal fluid (aCSF) containing (in mM) 125 NaCl, 2.5 KCl, 25 NaHCO_3_, 1.25 NaH_2_PO_4_, 2 CaCl_2_, 1 MgCl_2_, and 17 glucose (300 mOsm/kg). The brain was dissected, blocked, and transferred into a tissue-slicing chamber containing ice-cold ACSF. 250- to 300-μm-thick coronal slices containing LHb were cut using a vibratome. Following cutting, each slice was recovered individually in a pre-warmed (34°C) choline-based solution containing (in mM) 110 choline chloride, 11.6 ascorbic acid, 3.1 pyruvic acid, 2.5 KCl, 25 NaHCO_3_, 1.25 NaH_2_PO_4_, 0.5 CaCl_2_, 7 MgCl_2_, and 25 glucose. The time for recovery is determined by the age of the experiment animal and ranges from 10-30 min. Afterward recovery, slices were put in a secondary recovery chamber filled with 34°C ACSF for at least 20 min. The slices were then maintained in ACSF at room temperature until use. Choline and ACSF solutions were constantly bubbled with carbogen (95% O_2_ / 5% CO_2_).

### Ex vivo electrophysiology: measuring the sign of EP Sst→LHb synapses

For whole-cell recordings, individual slices were transferred to a recording chamber on the microscope. LHb neurons were visualized using an infrared differential interference contrast method under a 40× water-immersion Olympus objective. Epifluorescence (Excelitas, LED light source from X-Cite 120Q) was used to confirm virus expression and to identify regions displaying high density of axons from EP co-releasing neurons within the LHb. Recording pipettes (2–3 MΩ) were pulled from borosilicate glass using P-97 Flaming/Brown type micropipette puller (Sutter). Pipettes were filled with cesium-based internal recordings solution consisting of (in mM) 135 CsMeSO_3_, 10 HEPES, 1 EGTA, 4 Mg-ATP, 0.3 Na-GTP, 8 Na_2_- Phosphocreatine, 3.3 QX-314 (Cl- salt), pH adjusted to 7.3 with CsOH, and diluted to 290–295 mOsm/kg.

Whole-cell voltage-clamp recording was performed in acute slices continuously perfused with carbogenated ACSF at room temperature at a flow rate of 3∼4 ml/min. For all experiments, tetrodotoxin (TTX, 1 μM Tocris) and 4-Aminopyridine (4-AP, 400 μM Tocris) were added to the bath to ensure that monosynaptic currents from co-releasing EP neurons were measured. R,S-3- (2-carboxypiperazin-4- yl) propyl-1-phosphonic acid (CPP, 10 μM Tocris) was applied in bath solution to block NMDAR-mediated currents. Reported membrane potentials were not corrected for liquid junction potential (empirical=9.3 mV; theoretical=10.6 mV). After forming an intracellular seal with a target LHb neuron, 589 nm light stimulus (for ChrimsonR and ChRmine) or 473 nm (for ChR2) at 3-6 mW was delivered to the whole field-of-view to activate opsin-expressing EP Sst axons to confirm a synaptic transmission onto the postsynaptic cell. In LHb neurons that elicited PSCs, we first clamped the cell at -70 mV and delivered 5-10 stimulation pulses (2∼5 ms pulse duration, 10 s interstimulus interval, 1 pulse per sweep) to measure EPSCs before switching to 10 mV to measure IPSCs. In some cells, instead of clamping directly at 10 mV, we clamped the voltage from 0 mV and gradually increased the clamping voltage until no inward currents were observed.1 mg/ml biocytin was included in some cells to visualize the cell location in post-hoc histology.

Voltage-clamp recordings were amplified and low-pass filtered at 3 kHz using a Multiclamp700 B (Axon Instruments, Molecular Devices) and digitized at 10 kHz using an acquisition board (National Instruments). Data were saved with a custom version of ScanImage written in MATLAB.

### Immunohistochemistry: protocol

After each *ex vivo* experiment, brain slices were transferred to 4% paraformaldehyde and fixed overnight. Afterwards, slices were transferred to 1X phosphate-buffered saline (PBS) and stored in 4 °C until immunohistochemistry. Slices were mounted and imaged under 20X objective using a spinning-disk confocal microscope (Nikon).

For preparing brain slices after *in vivo* experiments, mice were euthanized and perfused transcardially with 1 M PBS followed by 4% paraformaldehyde (1 M). After 24 hour post-fix in 4% paraformaldehyde, brains were sliced into 60 µm thick coronal sections using a vibratome. Slices were mounted on glass slides with DAPI mounting medium (SouthernBiotech, 0100-20) and imaged under an automated slide scanner using a 10X objective (Olympus, VS200).

After brain slices were fixed, immunohistochemistry experiments were conducted as previously described (*47*). First, we washed the brain slices for 60 min in room temperature in 1% PBST solution (1X PBS and 0.1% TritonX-100). Second, we washed the slices in 10% blocking solution (10% Blocking One (Nacalai, 03953-95) in 0.3% PBST). Third, we added the primary antibody in the 10% blocking solution and let the slices sit overnight at 4 °C for slices after *in* vivo experiments and 3 days after *ex vivo* experiments. Fourth, we washed the slices in 0.3% PBST three times for 10 min each. Fifth, we incubated the slices in 10% blocking solution with the secondary antibody overnight at 4 °C. Sixth, we washed the slices in 0.3% PBST three times for 10 min each. Finally, we washed the slices in 1X PBS for at least 10 min, before mounting the slices.

### Immunohistochemistry: antibodies

The following antibodies were used with the source and dilution indicated in parentheses:

- Chicken anti-GFP (Abcam, ab13970; 1:2000)
- Rabbit anti-RFP (Rockland)
- Mouse anti-NeuN (Millipore Sigma, MAB377, 1:500)
- Streptavidin, Alexa Fluor 647 Conjugate (Invitrogen, S21374, 1:500)
- Donkey anti-chicken Alexa Fluor 488 (Jackson ImmunoResearch, 703-545-155, 1:1000)
- Donkey anti-rabbit Cy3 AffiniPure (Jackson ImmunoResearch, 711-165-152, 1:1000)
- Donkey anti-mouse Alexa Fluor 647-conjucated AffiniPure (Jackson ImmunoResearch, 715-605-151, 1:1000)

### Analysis: code

Most analysis in this paper are done in MATLAB. We create a neuroscience data analysis pipeline (NeuroDAP: https://github.com/shunnnli/NeuroDAP) with detailed documentation of most methods in the Wiki section of the repository. All the codes we used for data collection, processing, and analysis are contained inside.

### Analysis: syncing of in vivo recording

As mentioned in “*Behavior: behavior setup*”, syncing was done by sending an irregularly timed sync pulse to all recording systems. A custom MATLAB script was used to perform syncing.

Briefly, recordings of the sync pulse in different systems were extracted, and the onset of each sync pulse was identified. We then calculated the time interval between each sync pulse based on each system’s sampling frequency. The time interval of the sync pulse should be unique it each inter-pulse interval and common across all recording systems; thus, we can perform cross-correlation to extract the first common sync pulse. We defined the time of the first common sync pulse as the official start of the recording sessions. In practice, all systems were turned on within one minute.

After identifying the session start time, we use the recording system with the highest sampling rate (normally recorded by NIDAQ in SpikeGLX, which is 10 kHz) to interpolate a basis time vector based on sampling rate. In this vector, each element indicates when that sample was acquired relative to the session start. Since the sync pulses are sent to all systems, for systems with slower sampling rate, we used our basis time vector to first locate the time of all sync pulse onsets, then interpolate the time based on sampling frequencies of the corresponding system. For more detailed explanation on the algorithm, please refer to the Wiki section of NeuroDAP repository.

### Analysis: fiber photometry processing

In general, all photometry recordings are demodulated and downsampled to 50 Hz. The frequency modulated signals were detrended using a rolling z-score with a time window of 3 minutes. As the ligand-dependent changes in fluorescence measured *in vivo* are small (few %) and the frequency modulation is large (∼100%), the variance in the frequency modulated signal is largely ligand independent. Thus, z-scoring on a large time window eliminates photobleaching without affecting signal. Detrended, frequency-modulated signals were frequency demodulated by calculating a spectrogram with 1 Hz steps centered on the signal carrier frequency using the MATLAB’s spectrogram() function. The spectrogram was calculated in windows of 80 samples with 40 sample overlap, corresponding to a final sampling frequency of 50 Hz. The demodulated signal was calculated as the power averaged across a 3 Hz frequency band centered on the carrier frequency. The demodulated signal was passed through an additional rolling z-score (180 sec window) to quantify the fluorescence transients as z-scores. In some sessions, the dLight recording is modulated at 171 Hz. In those recordings, the spectrogram was calculated in windows of 216 samples with 196 sample overlap. The demodulated signal was calculated as the power averaged across a 5 Hz frequency band centered on the carrier frequency. For Fig. 1, demodulated data were then low-pass filtered at 8 Hz to remove high-frequencies oscillations due to imperfect alignment of frequency demodulation and then underwent the additional z-score window afterwards. Data were then resampled to 50 Hz using MATLAB’s rat() and resample() function. See demodulateSignal() in NeuroDAP repository for more details and code.

### Analysis: behavior and photometry

For all recording sessions, a trial table was generated to assign all behavior events (e.g. licking, optogenetic stimulations) to their respective trials. For baseline recording sessions, each trial was separated into two stages. One is 0-2 s since the start of a trial (stimuli delivery), and second is 2s to the start of the next trial. For reward and punish pairing sessions, each trial was separated into three stages, corresponding to the cue, response, and free outcome period. For analyzing the effect of WiChR-mediated optogenetic inhibition of LHb, we examined dopamine responses up to 1 s after the stimulation onset.

For licking, we defined anticipatory licks as lick events that precede outcome delivery. For photometry, at each stage the maximum, minimum, and amplitude of recorded transients were calculated. The amplitude of the photometry transient (ΔDA or ΔGlu) was defined as:

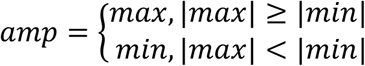

where max and min indicates the maximum or minimum of transients during the defined window for each stage.

To reveal changes over trials due to learning of as reflected in each metric (e.g. dopamine amplitude or number of anticipatory licking), we first extracted all metrics for every trial and calculated averages in bins of 10 trials (e.g. **Fig. 1G**). We also calculated the slope of each metric vs. trial number using the grouped data. Since each trial type has different probability of occurrence throughout the session (e.g. 50% for opto-only and 40% for opto+tone trials), we normalized the trial number to make it in range [0, 1] and calculated the slope relative to this normalized range. Such normalization essentially converted the x axis from a trial-unit frame to a time-unit frame.

### Analysis: ex vivo electrophysiology data processing

In each acquisition, a -5 mV pulse lasting 100 ms is delivered to measure passive cellular and recording parameters (i.e. RC check). Based on the location of optogenetic stimulation and RC check, we divided each sweep into following windows:

1. Baseline window: data before stimulation onset and data between 200 ms after stimulation onset and before RC check onset.
2. Control window: data 30 ms before stimulation onset.
3. Analysis window: data 30 ms after stimulation onset.
4. Peak window: data 1 ms around the maximum or minimum data point during analysis window for IPSC and EPSC, respectively.

We then calculated the baseline-subtracted version of the recorded data. Baseline is calculated by averaging all data recorded during the baseline window, which is subtracted from the raw recordings to generate a baseline-corrected recoding for each sweep. For both the baseline and analysis window, we find the time of maximum and minimum values and average data points collected 2 ms around this maximal response window. To calculate the amplitude of EPSC responses, the minimal current response during the 2 ms response window was used to calculate the EPSC amplitude and the opposite is true for amplitudes of IPSC responses. Peak response during the baseline window of the same direction (minimum for EPSC, maximum for IPSC) was used as control. To calculate total charges of EPSC or IPSC responses, the sum of recorded currents during the entire 30 ms analysis window was used.

After dividing sweeps into these windows, we calculate quality metrics of the cell using the RC check. We only include data collected with stable recordings. We use following quality metrics to select cells and inspect each cell manually after initial quality checks. Series resistance (Rs) was computed for every trial by fitting an RC response curve with an exponential decay and extrapolating back to the instantaneous peak size at the start of the voltage-pulse. We eliminated sweeps in which Rs of the cell was more than 30 MΩ and the whole trials if there are less than 5 qualified sweeps. These cells are small with generally high input resistance (mean Rm across recorded LHb neurons is 478.69 MΩ), permitting use of recordings with higher Rs than the typical 20 MΩ cut off. We also excluded cells for which the opto-evoked responses did not differ from baseline current fluctuations, indicating a lack of a synapse between the stimulated axons and the recorded voltage-clamped neuron. Sweeps with large voltage error (> 10 mV) and large voltage drifts were also eliminated. No relationships between synaptic sign index and recording properties were observed (**fig. S10**) and passive cellular and recording properties did not differ across conditions. See loadSlices() and plotEpochSummary() in NeuroDAP for corresponding code implementation.

### Analysis: Synaptic sign index

Synaptic sign index is essentially the difference between excitatory and inhibitory currents amplitudes normalized by the total amount of current received. It is calculated by the following formula:

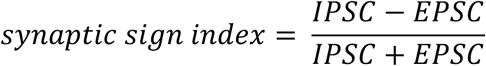

where EPSC and IPSC here represents the amplitude of excitatory and inhibitory currents, respectively (see above for details of amplitude calculation). We also calculated an analogous version of synaptic sign index for each cell using the total charge of the cell’s synaptic currents instead of amplitude.

### Analysis: correlating synaptic sign index with dopamine updates

Early and late window of each behavioral session are defined by the first n trials or the last n trials, respectively, of the final recording session before electrophysiological analysis. To calculate the slope of changes in DA transient amplitudes within each window, we first performed a moving average of DA amplitude during cue period of opto-only trials across the nearby five trials of the same type to reduce noise due to single-trial variations. This will generate two vectors of size (#animals, 1). They represent dopamine amplitude slope across trials during early or late window for each animal, respectively. For **Fig. 4**, we also calculated the average DA amplitude evoked by EP opto stim across the last session for each animal, generating a third vector of the same size as above. For **fig. S14**, we calculate the average DA amplitude evoked by EP opto stim across the selected trial window length. We aligned the trials from the start or from the end to ensure we did not observe spurious correlations. These DA-related vectors are then correlated with animal sign index, which is calculated by averaging the synaptic sign index of all the cells recorded from that animal.

Spearman’s correlation coefficient and slope fit were calculated to assess the correlation. For slope fit, we performed a linear fit weighted by the number of cells recorded from each animal. This is because animals with more cells recorded are more likely to represent the true underlying distribution of synaptic sign index across all EP Sst→LHb synapses. Spearman’s correlation coefficient as calculated using MATLAB’s corr() function.

### Analysis: reclassifying recorded cells based on recent dopamine updates

Based on results in **fig. S14**, we found that dopamine amplitude slopes calculated from last 40 to 15 trials in the session correlates with EP Sst→LHb synaptic sign index. Therefore, we averaged the DA amplitude slopes calculated from last 40 to last 15 trials for each animal. We then performed K-means clustering using MATLAB with 100 replicates to classify animals into three groups, representing increasing, stable, and decreasing DA amplitudes during the above window.

We also separated data into training and test sets to avoid misidentifying spurious correlation using the whole dataset. We randomly separated all animals in the dataset into training (80% of animals) and test (20% of animals) sets. Using data recorded from the training set animals, we performed the same sweeping trial window analysis as above (see section above) to identify trial stretches with significantly correlated animal sign index and DA slope of interest (early or late DA slope, average DA amplitude aligned to start or end of the session). If no trial stretches are found to be significantly correlated, we use the whole session to perform our analysis below.

These trial stretches with correlated animal sign index and DA slope found in training set were then used to calculate the correlation between animal sign index and DA slope in animals from the test set. We also calculated the average synaptic sign index and cumulative probability of synaptic sign index from cells of test animals based on their respective DA slope group (increasing/stable/decreasing DA). We repeated this process for 200 times.

### Analysis: statistics

All statistical analyses were performed using custom code written in MATLAB. All analysis of statistical comparisons in the first 2 figures and their supplementary figures were done using Kolmogorov–Smirnov tests unless stated otherwise. For calculating significance of synaptic sign index across different conditions, we used bootstrapping. Specifically, we pooled all the cells together and randomly selected cells with the same sample size of the original condition. We then calculated the mean synaptic sign index from the bootstrapped group and repeat such process for 10000 times. The significance of Spearman correlation coefficients was calculated by MATLAB’s corr() function. No statistical methods were used to predetermine sample sizes, but our sample sizes are similar to those reported in previous publications. The corresponding p value range relating to the number of asterisks in the figure are as follows:

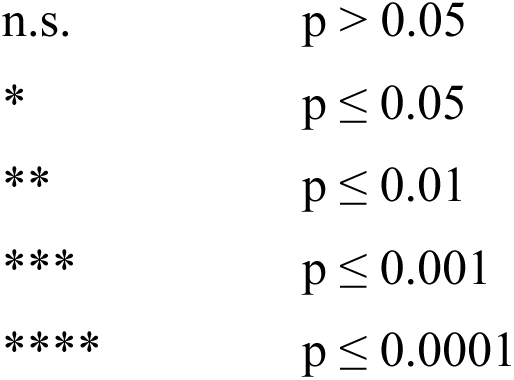

**Fig. S1.**
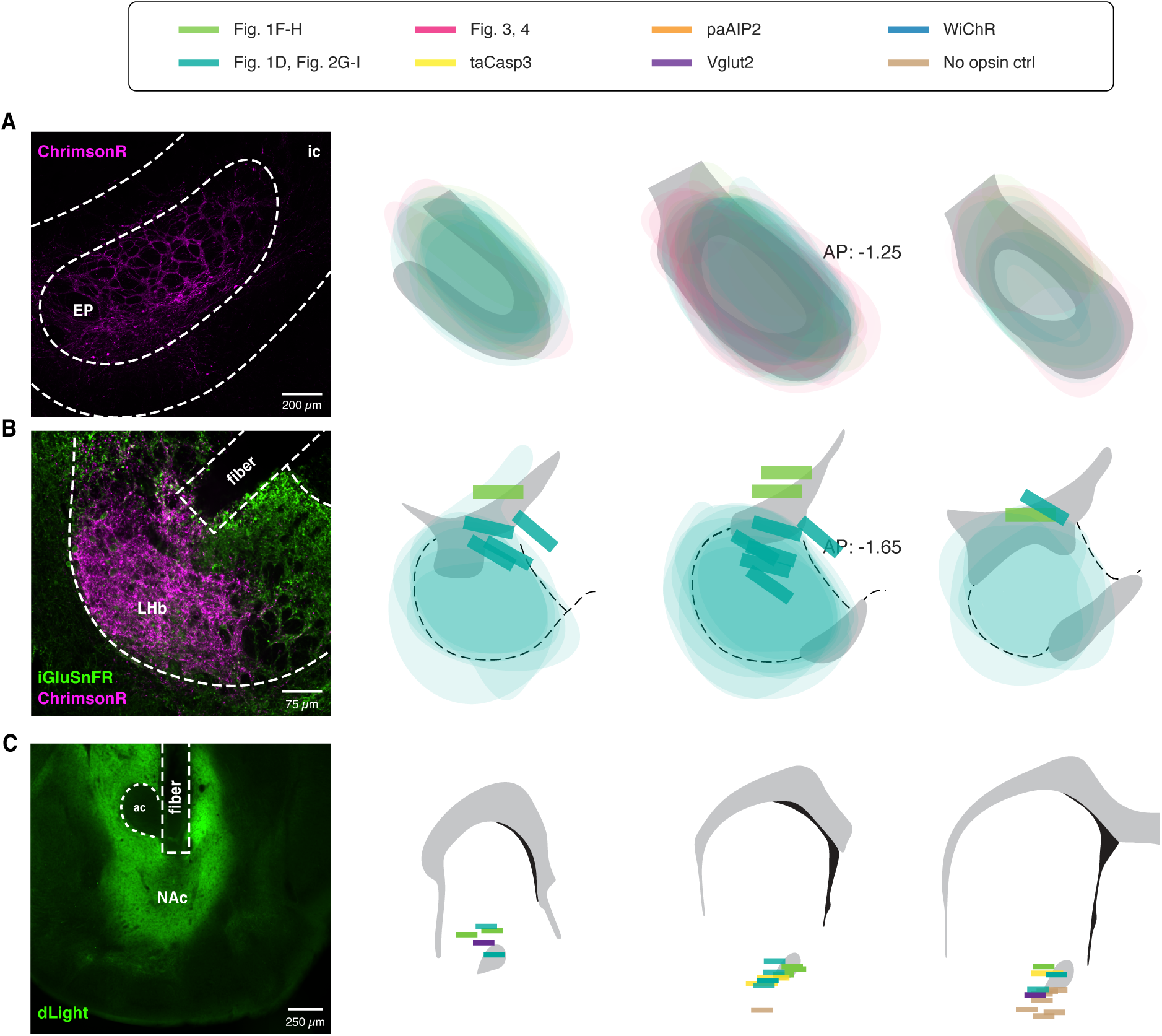
Expression maps and example histology of EP, LHb, and NAc. **A**, Example expression of ChrimsonR in EP (*left*) with expression map of ChrimsonR in animals included in **Fig. 1**, **Fig. 2G-I** (animals with iGluSnFR expressed in LHb), and **Fig. 3, 4** (*right*). **B**, Example expression of iGluSnFR and ChrimsonR in LHb and implanted optical fiber (*left*) with expression map of iGluSnFR in LHb neurons and approximate location of fiber placement in the same animals as **a** (*right*). **C**, Example expression of dLight in NAc and implanted optical fiber (*left*) with expression map of dLight and fiber placement in subsets of animals with recovered NAc. Colors are sorted by experiment cohort (indicated at the top, *right*).

**Fig. S2.**
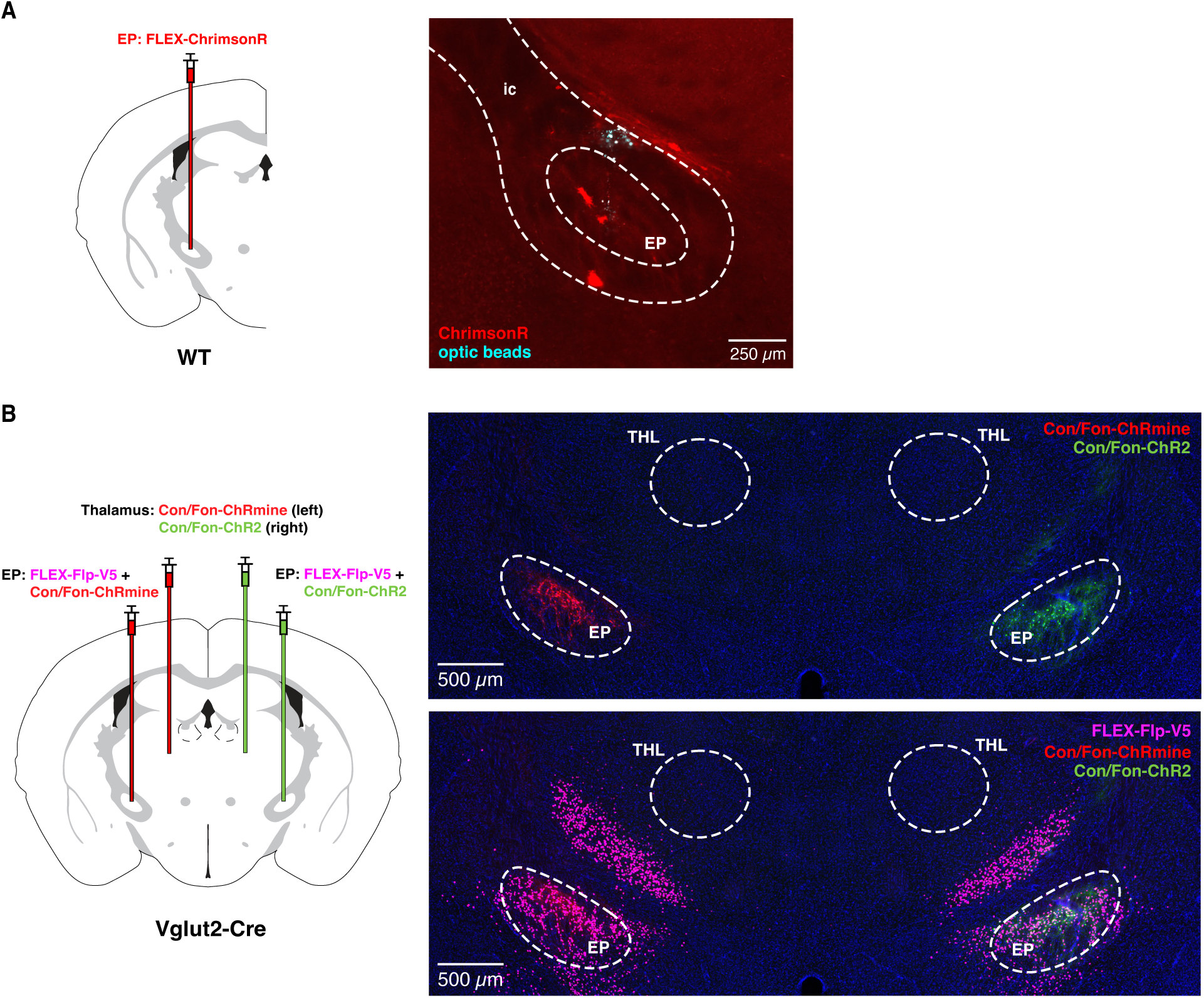
Expression of excitatory opsins used in EP are recombinase dependent. **A**, Schematics of viral transduction to validate Cre specificity of FLEX-ChrimsonR (*left*) and corresponding histology (*right*). Virus was injected into EP of WT animal along with fluorescent optic beads (cyan) to mark the injection site. As seen in the histology, other than one or two cells expressed ChrimsonR in ventral TRN (which do not project to LHb, see **fig. S5**), there is no obvious expression of ChrimsonR in EP (we think the three red dots are tissue debris and not ChrimsonR-expressing neurons). **B**, Schematics of viral transduction to validate Cre and Flp specificity of Con/Fon-ChR2 and Con/Fon-ChRmine (*left*) and histology (*right*). Viruses are injected into thalamus (THL) in Vglut2-Cre animals, where no opsin expression are observed in thalamus. As a positive control, we co-injected FLEX-Flp-V5 with ChR2 or ChRmine viruses in EP. This leads to significant opsin expression in EP for both ChR2 and ChRmine. The two histology images on the right are of the same samples with or without showing fluorescent channel containing labeling of the V5 tag from the FLEX-Flp-V5 virus.

**Fig. S3.**
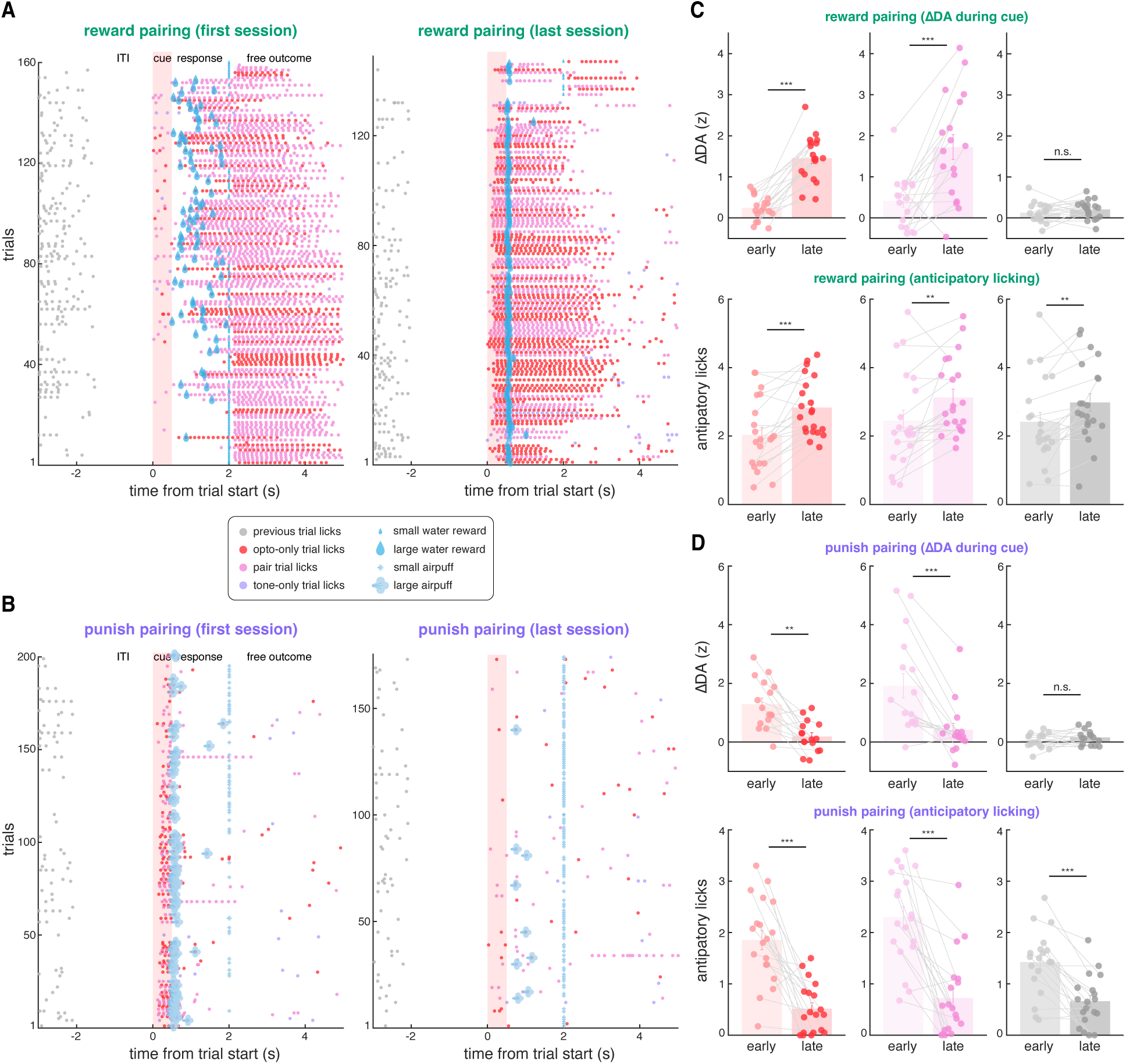
Animals learns pairing between EP opto stim and reward/aversive outcomes. **A,** Lick raster plot of an example animal during the first (*left*) and last (*right*) session of reward pairing. Red and pink dots indicate lick events during opto-only trials and opto+tone trials, respectively. The timing of reward delivery at each trial are indicated using the water icon. Animal gradually learned cue-reward pairing as seen by increased number of licks during cue and response period and increase likelihood to obtain large water reward. **B**, Lick raster plot of the same animal during the first (*left*) and last (*right*) session of punishment pairing (after finishing sessions of reward pairing in **A**). Red and pink dots indicate lick events during opto-only trials and opto+tone trials, respectively. The timing of air puff delivery at each trial are indicated using the air puff icon. Animal gradually learned cue-punishment pairing as seen by decreased number of licks during cue and response period and increase likelihood to avoid large air puff punishment. **C**, The averaged amplitude of cue-evoked DA transients (*top*) and amount of anticipatory licking (*bottom*) during early or late phase (first or last 40 trials) of the reward pairing. Data from combining animals in **Fig. 1F-H** and **Fig. 2G-I**. (*top*: n=18 mice, *bottom*: n=21 mice). Opto-only trials, opto+tone trials, and randomly selected control timepoints are plotted in red, pink, and gray, respectively. Each dot is the averaged amplitude of DA transients or anticipatory licks for an animal, average ± s.e.m are also shown. **D**, Same as **C** but data for punishment pairing (combining animals in **Fig. 1F-H** and **Fig. 2G-I**, *top*: n=16 mice, *bottom*: n=19 mice).

**Fig. S4.**
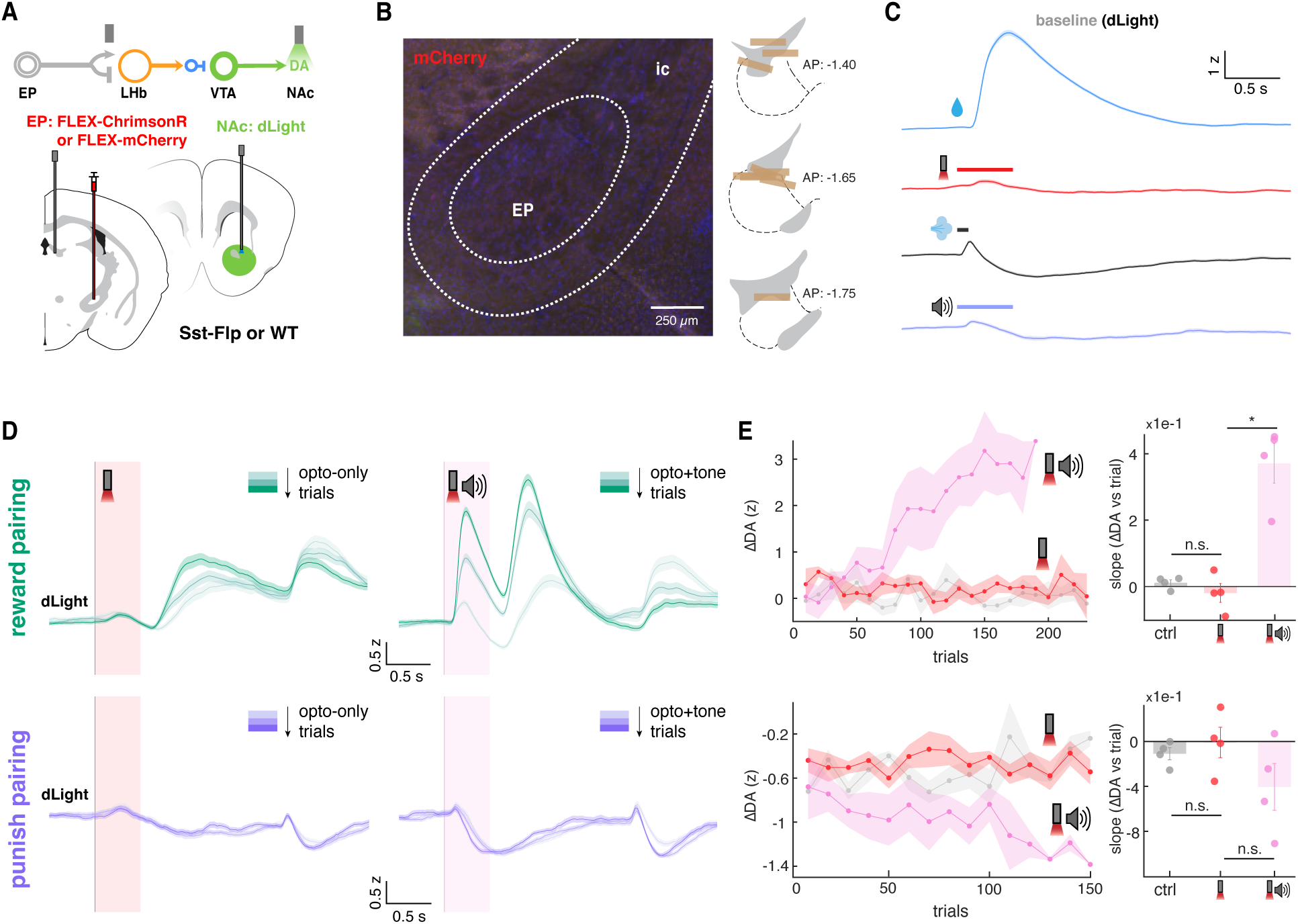
Animals with no opsin expression in EP fail to learn opto-outcome pairing. **A**, Schematic of viral transductions and photometric recordings. Cre-dependent ChrimsonR or mCherry is injected into EP of Sst-Flp mice, resulted in the lack of opsin expression in EP Sst neurons. An optical fiber is implanted in LHb to deliver red light into LHb to mimic normal optogenetic stimulation. Fluorescent dopamine sensor dLight is expressed in NAc to monitor DA release through a second optical fiber. **B**, Example expression of mCherry in EP (*left*) and approximate location of recovered implanted optical fibers in LHb used in this experiment (*right*, n=8 mice). **C**, NAc DA transients aligned to onset of water reward (blue), opto stim-mimicking light pulses (red), air puff (black), and auditory tone (purple) during baseline sessions (n=8 mice). The average z-scored signals across trials from all animals ± s.e.m. are shown. **D**, NAc DA transients aligned to onset of opto-only cue (left, red) or opto+tone cue (right, pink) during reward pairing sessions (*top*) and punishment pairing sessions (*bottom*) (n=4 mice each for reward or punishment pairing). Trials are color-coded from light to darker color indicating early to late trials (each color is around 200 trials across all animals, i.e. 50 trials for each animal). The average z-scored signals across trials from all animals ± s.e.m. are shown. **E**, The amplitude of cue-evoked NAc DA transients (ΔDA) as a function of trial number (*left*) and corresponding slope between changes in evoked DA transients and trial number (*right*, ΔDA vs trials). Reward pairing data are plotted on the top and punishment pairing are plotted at the bottom. Data are shown for trials with the opto-only cue (red), opto+tone cue (pink), or randomly selected recording timepoints throughout each session as control (gray). On the left, each dot indicates the average DA amplitudes of the nearby 10 trials and the averaged z-scored amplitudes across animal ± s.e.m. are shown. On the right, the data for individual mice (dots) and the average ± s.e.m are shown.

**Fig. S5.**
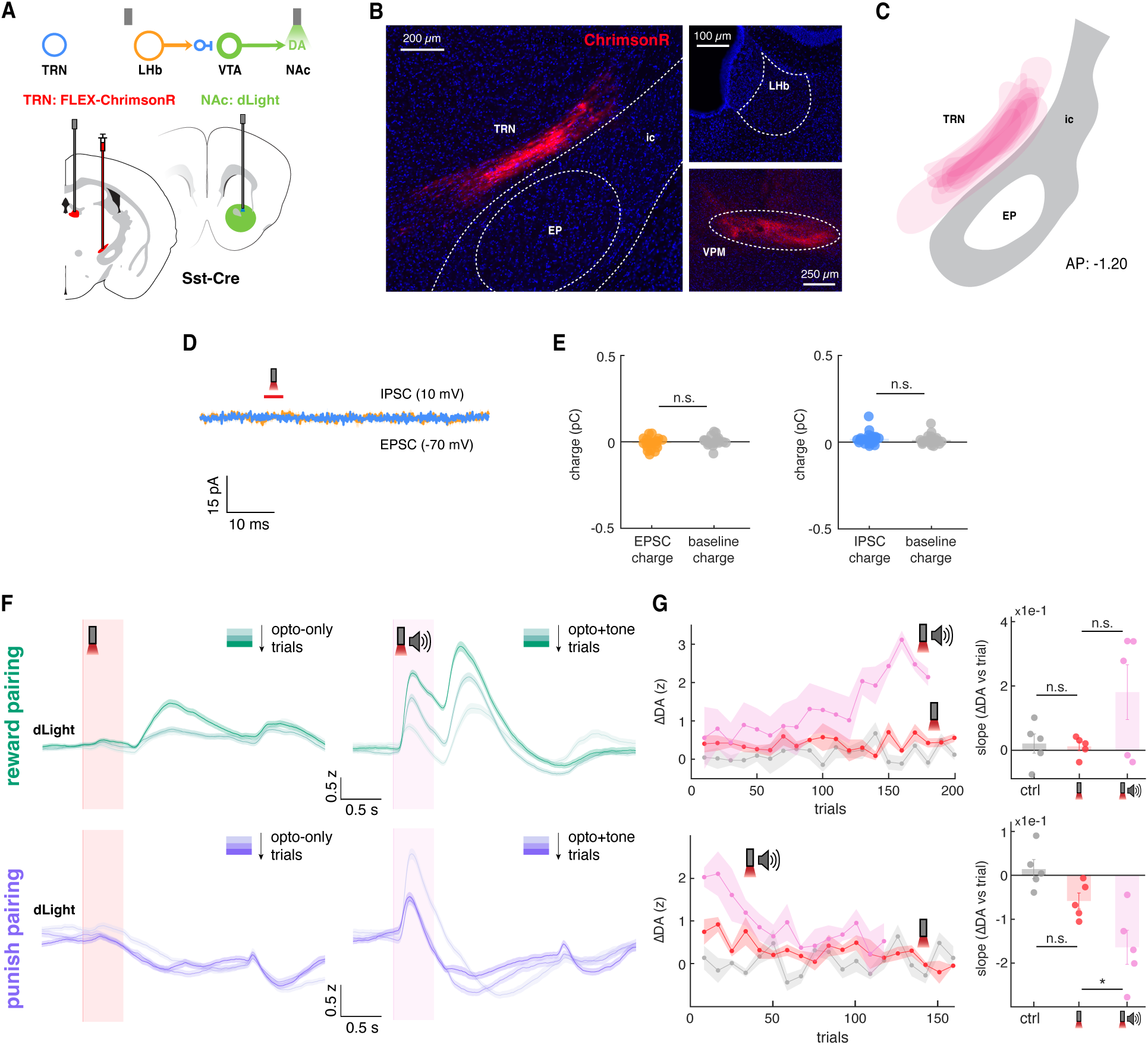
Ventral TRN neurons do not project to LHb and animals with opsin expressed in TRN fail to learn opto-outcome pairing. **A**, Schematic of the viral transductions and photometric recordings. Cre-dependent ChrimsonR is expressed in ventral TRN in Sst-Cre mice. An optical fiber is implanted in LHb to deliver red light into LHb for optogenetic stimulation. Fluorescent dopamine sensor dLight is expressed in NAc to monitor DA release through a second optical fiber. **B**, Example expression of ChrimsonR in ventral TRN (*left*) and its putative downstream targets (*right*). Note that while there is apparent expression of ChrimsonR-positive fibers around ventral posteromedial nucleus (VPM) of thalamus, no expression of ChrimsonR-positive fibers is observed in LHb, indicating a lack of projection. **C**, Expression map of ChrimsonR virus in ventral TRN. **D**, Example post-synaptic currents (PSC) of a LHb neuron recorded at -70 mV (yellow) and 10 mV (blue). Onset of optogenetic stimulation of putative ventral TRN ChrimsonR-expressing fibers in LHb are indicated in red. **E**, Total charge of EPSCs (*left*) and IPSCs (*right*) in LHb neurons (n=23 cells, N=4 mice) comparing to charge of spontaneous events during baseline periods. **F**, NAc DA transients aligned to onset of opto-only cue (left, red) or opto+tone cue (right, pink) during reward pairing sessions (*top*) and punishment pairing sessions (*bottom*) (n=5 mice). Trials are color-coded from light to darker color indicating early to late trials (each color is around 200 trials across all animals, i.e. 50 trials for each animal). The average z-scored signals across trials from all animals ± s.e.m. are shown. **G**, The amplitude of cue-evoked NAc DA transients (ΔDA) as a function of trial number (*left*) and corresponding slope between changes in evoked DA transients and trial number (*right*, ΔDA vs trials). Reward pairing data are plotted on the top and punishment pairing are plotted at the bottom. Data are shown for trials with the opto-only cue (red), opto+tone cue (pink), or randomly selected recording timepoints throughout each session as control (gray). On the left, each dot indicates the average DA amplitudes of the nearby 10 trials and the averaged z-scored amplitudes across animal ± s.e.m. are shown. On the right, the data for individual mice (dots) and the average ± s.e.m are shown.

**Fig. S6.**
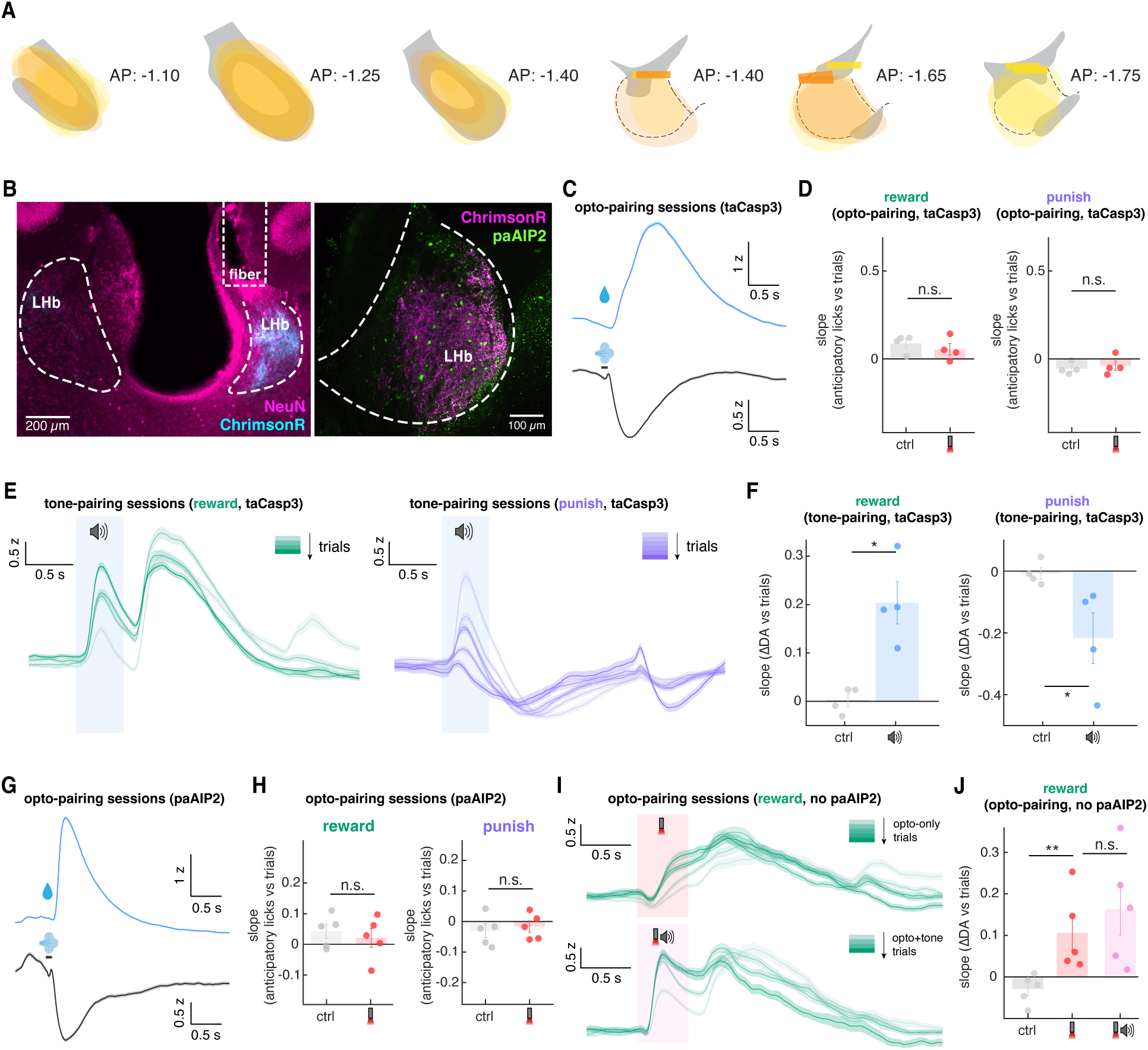
Control experiments for taCasp3 and paAIP2 cohorts. **A**, *Left*, expression map of Cre-dependent ChrimsonR expression in EP. *Right*, expression map of taCasp3 and paAIP2, as well as recovered optical fiber locations in LHb (yellow is taCasp3 cohort, orange is paAIP2 cohort). **B**, *Left*, example histology of taCasp3-mediated LHb lesion. ChrimsonR-expressing fibers from EP and optical fiber location are also included. Note that taCasp3-expressed LHb (right in the picture), comparing to the uninjected side, have significantly smaller area and less intact neurons based on NeuN staining (the contour of LHb on the right is approximate due to leison). The general higher level of NeuN fluorescent on the injected/implanted side is to be expected because the use of mouse antibody. ChrimsonR-expressing EP axons are not affected by caspase expression in LHb. *Right*, example histology showing ChrimsonR-expressing EP axons and paAIP2 expression in LHb. **C**, NAc DA transients as reported by dLight fluorescence aligned to onset of water delivery (blue) or air puff (black) during opto-pairing sessions for taCasp3 cohort. The average z-scored signals ± s.e.m. are shown. Note that dopamine increases to water reward and decreases to air puff, suggesting that impairment in learning is not caused by animals’ inability to detect reward or punishment stimuli. **D**, Slope between changes in anticipatory licking and trial number for reward pairing (*left*) and punishment pairing (*right*). Data are shown for trials with the opto-only cue (red) or randomly selected recording timepoints throughout each session as control (gray). The data for individual mice (dots) and the average ± s.e.m are shown. **E**, NAc DA transients aligned to onset of tone (blue) during reward tone-pairing sessions (*left*) and punishment tone-pairing sessions (*right*) (same mice from taCasp3 cohort as in Fig 2). Trials are color-coded from light to darker color indicating early to late trials (each color is around 200 trials across all animals, i.e. 50 trials for each animal). The average z-scored signals across trials from all animals ± s.e.m. are shown. **F**, Slope between changes in evoked DA transients (ΔDA) and trial number for reward tone- pairing (*left*) and punishment tone-pairing (*right*). Data are shown for trials with the tone-pairing cue (blue) or randomly selected recording timepoints throughout each session as control (gray). The data for individual mice (dots) and the average ± s.e.m are shown. **G**, NAc DA transients as reported by dLight fluorescence aligned to onset of water delivery (blue) or air puff (black) during opto-pairing sessions for paAIP2 cohort. The average z-scored signals ± s.e.m. are shown. Note that dopamine increases to water reward and decreases to air puff, suggesting that impairment in learning is not caused by inability to detect reward or punishment stimuli. **H**, Slope between changes in anticipatory licking and trial number for reward opto-pairing (*left*) and punishment opto-pairing (*right*). Data are shown for trials with the opto-only cue (red) or randomly selected recording timepoints throughout each session as control (gray). The data for individual mice (dots) and the average ± s.e.m are shown. **I,** NAc DA transients aligned to onset of opto-only cue (top, red) or opto+tone cue (bottom, pink) during reward opto-pairing sessions without paAIP2-mediated CaMKII inhibition (same mice from paAIP2 cohort as in **Fig. 2D-F**). Trials are color-coded from light to darker color indicating early to late trials (each color is around 200 trials across all animals, i.e. 50 trials for each animal). The average z-scored signals across trials from all animals ± s.e.m. are shown. **J**, Slope between changes in evoked DA transients and trial number for reward opto-pairing without paAIP2 inhibition. Data are shown for trials with the opto-only cue (red), opto+tone cue (pink), or randomly selected recording timepoints throughout each session as control (gray). The data for individual mice (dots) and the average ± s.e.m are shown.

**Fig. S7.**
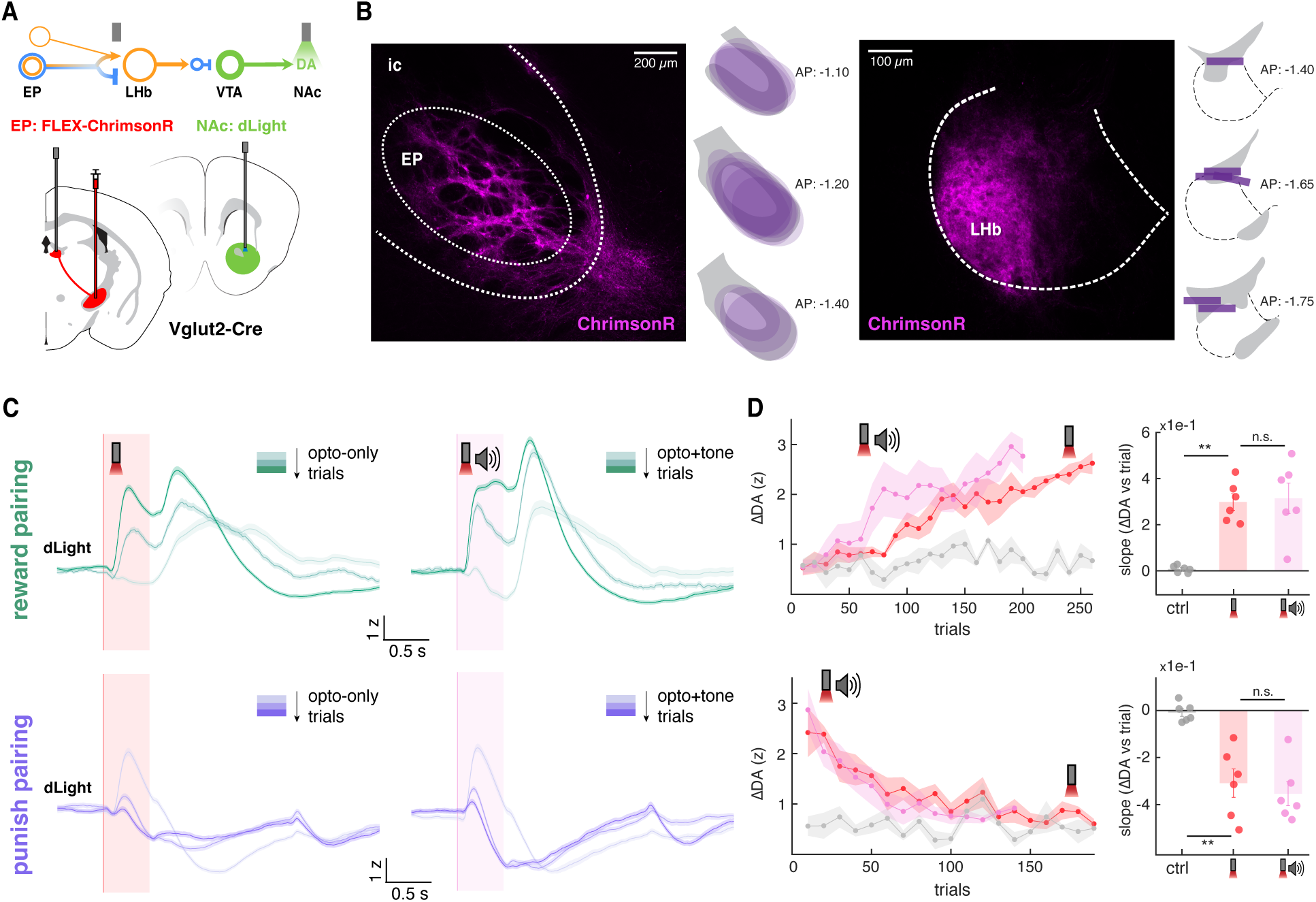
Vglut2-Cre animals also learned opto-outcome pairing. **A**, Schematic of viral transductions and photometric recordings for LHb Vglut2 experiment. Cre-dependent ChrimsonR is selectively expressed in EP of *Vglut2-Cre* mice. This strategy labels the Sst+ co-releasing neurons and the purely glutamatergic PV+ neurons that also solely project to LHb. An optical fiber is implanted in LHb to optogenetically stimulate EP ChrimsonR- expressing axons in LHb. dLight is expressed in NAc to monitor DA release through a second optical fiber. **B**, *Left*, example histology of Cre-dependent ChrimsonR expression in EP with expression map in EP across all animals in experiment cohort. *Right*, example histology of ChrimsonR- expressing EP axons in LHb with approximate optical fiber locations in LHb across all animals in experiment cohort. **C**, NAc DA transients aligned to onset of opto-only cue (left, red) or opto+tone cue (right, pink) during reward pairing sessions (*top*) and punishment pairing sessions (*bottom*) (n=6 mice). Trials are color-coded from light to darker color indicating early to late trials (each color is around 200 trials across all animals, i.e. 50 trials for each animal). The average z-scored signals across trials from all animals ± s.e.m. are shown. **D**, The amplitude of cue-evoked NAc DA transients (ΔDA) as a function of trial number (*left*) and corresponding slope between changes in evoked DA transients and trial number (*right*, ΔDA vs trials). Reward pairing data are plotted on the top and punishment pairing are plotted at the bottom. Data are shown for trials with the opto-only cue (red), opto+tone cue (pink), or randomly selected recording timepoints throughout each session as control (gray). On the left, each dot indicates the average DA amplitudes of the nearby 10 trials and the averaged z-scored amplitudes across animal ± s.e.m. are shown. On the right, the data for individual mice (dots) and the average ± s.e.m are shown.

**Fig. S8.**
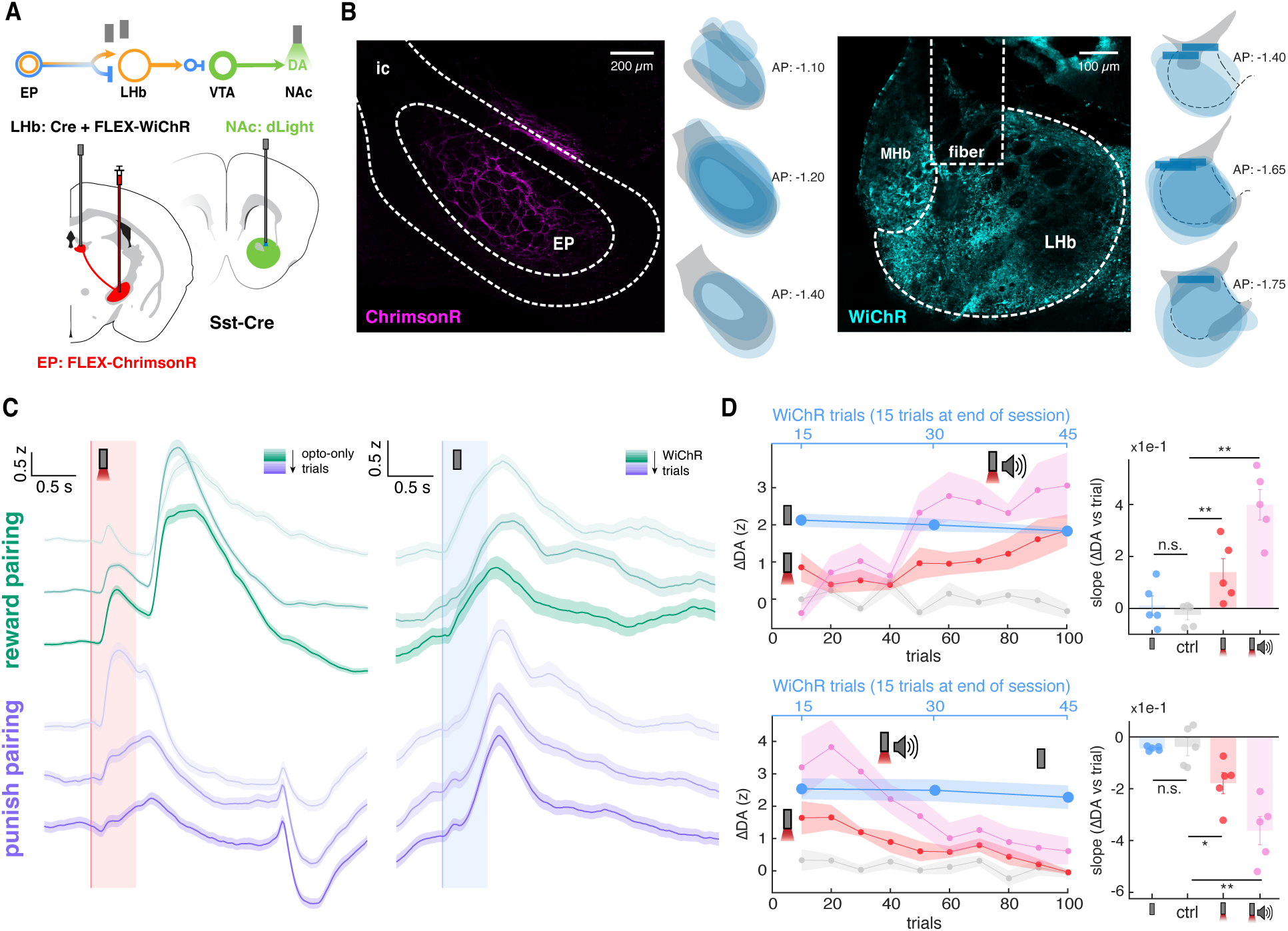
LHb always inhibit dopamine throughout opto-outcome pairing. **A**, Schematic of viral transductions and photometric recordings for LHb WiChR experiment. Cre-dependent ChrimsonR is selectively expressed in EP Sst neuron of *Sst-Cre* mice. An optical fiber is implanted in LHb to optogenetically stimulate EP ChrimsonR-expressing axons in LHb. dLight is expressed in NAc to monitor DA release through a second optical fiber. FLEX-WiChR and Cre are co-expressed in LHb to inhibit LHb neurons. WiChR are stimulated for 15 times after the conclusion of every pairing session to access the effect of LHb inhibition on NAc DA release. **B**, *Left*, example histology of Cre-dependent ChrimsonR expression in EP with expression map in EP across all animals in experiment cohort. *Right*, example histology of WiChR expression in LHb with the corresponding expression map and approximate optical fiber locations in LHb across all animals in experiment cohort. **C**, NAc DA transients aligned to onset of opto-only cue (*left*, red) and WiChR stimulation after each session (*right*, blue) during reward pairing sessions (green) and punishment pairing sessions (purple) (n=5 mice). Trials are color-coded from light to darker color indicating early to late trials (Opto-only cue aligned traces: each color is around 200 trials across all animals, i.e. 50 trials for each animal; WiChR aligned traces: each color is 45 trials across all animals, i.e. 15 trials for each animal). The average z-scored signals across trials from all animals ± s.e.m. are shown. **D**, The amplitude of evoked NAc DA transients (ΔDA) as a function of trial number (*left*) and corresponding slope between changes in evoked DA transients and trial number (*right*, ΔDA vs trials). Reward pairing data are plotted on the top and punishment pairing are plotted at the bottom. Data are shown for trials with the opto-only cue (red), opto+tone cue (pink), WiChR- mediated LHb inhibition (blue), or randomly selected recording timepoints throughout each session as control (gray). On the left, each dot indicates the average DA amplitudes of the nearby 10 trials (15 trials for WiChR) and the averaged z-scored amplitudes across animal ± s.e.m. are shown. On the right, the data for individual mice (dots) and the average ± s.e.m are shown.

**Fig. S9.**
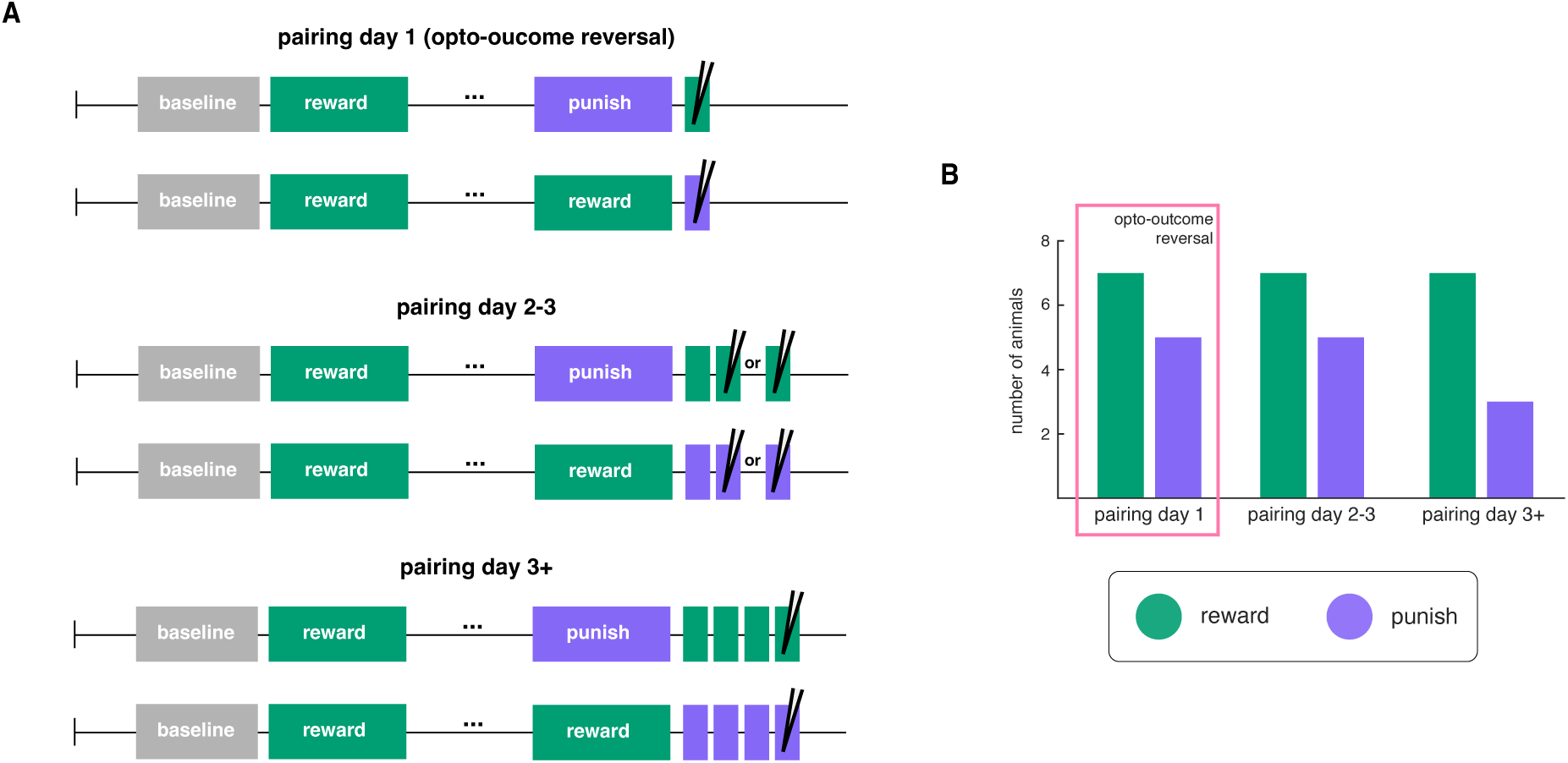
Experiment schedules for *in vivo* training to *ex vivo* electrophysiology. **A**, Timeline of three types of experiment schedules. All animals will undergo baseline and reward pairing phase, so that we can ensure all animals that undergo dissection and *ex vivo* recording are able to successfully learn the opto-outcome association. For pairing day 1 cohorts, they will end with a full punishment or reward pairing phase, followed by one day of reward or punishment pairing, respectively (i.e. opto-outcome reversal). *ex vivo* experiments are conducted immediately after the reversal session. For pairing day 2-3 cohorts, after a full punishment or reward pairing phase, the animals go through two or three days of reward or punishment pairing sessions, respectively (i.e. 2-3 sessions after opto-outcome reversal). *ex vivo* experiments are conducted after the second or third sessions. Similarly, pairing day 3+ cohorts go through 3+ opto-outcome pairing sessions after opto-outcome reversal before *ex vivo* recording. **B**, Number of animals used for each type of experiment schedules. Cells recorded at pairing day 1 (i.e. opto-outcome reversal) are plotted in **Fig. 3**.

**Fig. S10.**
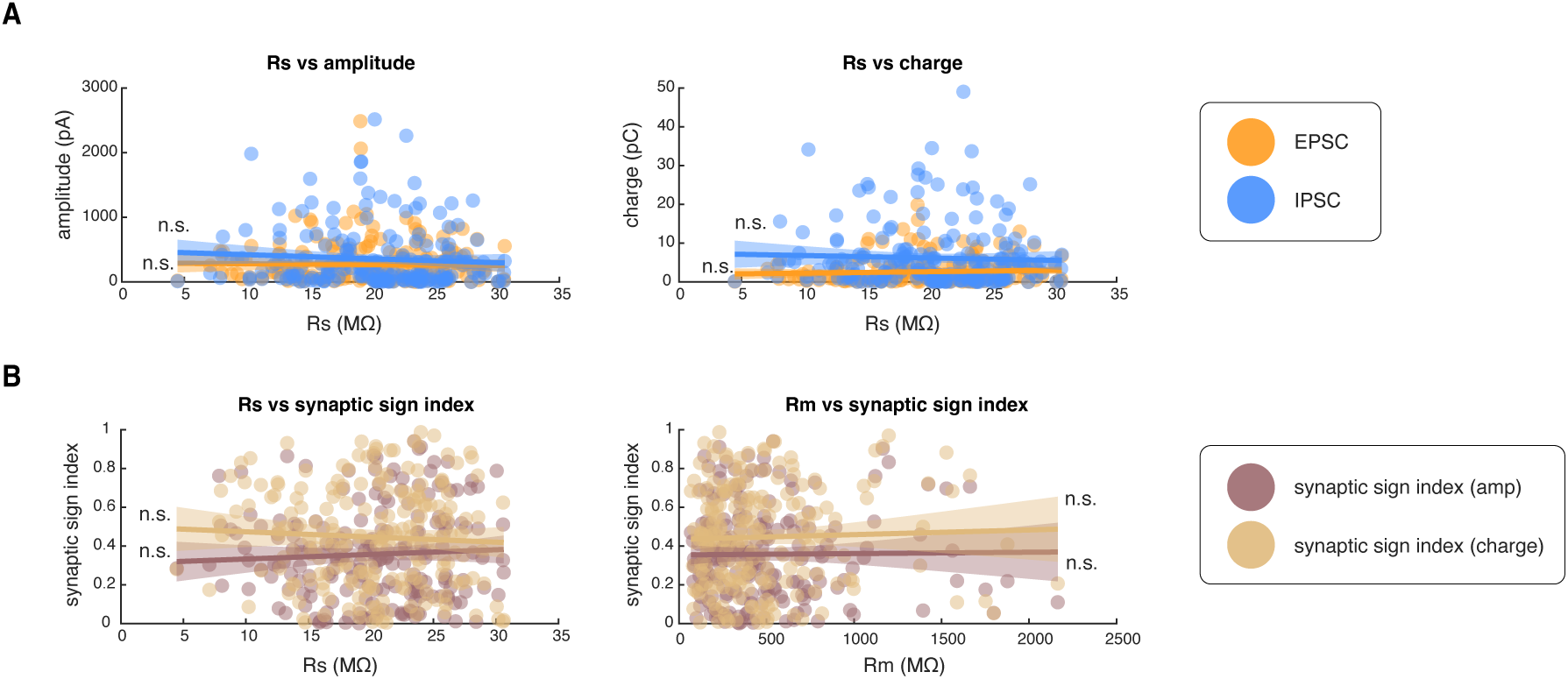
Synaptic sign index of each cell versus recording parameters. **A**, Recorded peak amplitude (*left*) and total charge (*right*) of excitatory and inhibitory postsynaptic currents (EPSC: yellow, IPSC: blue) as a function of series resistance. Solid line shows the best fit line and shaded region correspond to the 95% confidence interval. Significances are calculated by bootstrapping. **B**, Synaptic sign index calculated based on amplitude (brown) and total charge (beige) as a function of series resistance (*left*) and membrane resistance (*right*). Solid line shows the best fit line and shaded region correspond to the 95% confidence interval. Significances are calculated by bootstrapping.

**Fig. S11.**
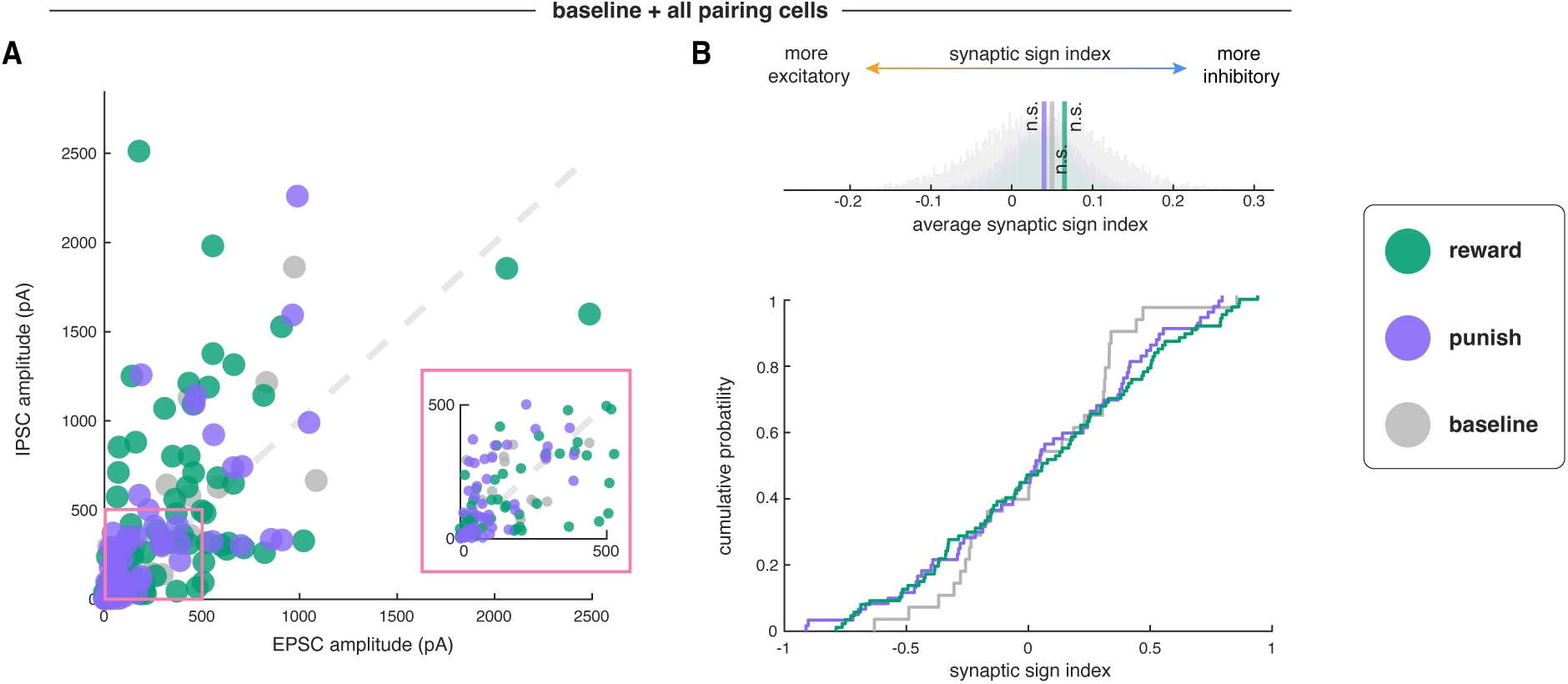
Synaptic signs of EP Sst**→**LHb synapses do not differ when classifying cells based on opto-outcome pairing direction across all conditions. **A**, Scatter plot of EPSC amplitude and IPSC amplitude of all cells recorded. Each dot represents a cell. Cells are colored based on whether EP opto stim are paired with reward (green, n=88 cells, 21 mice) or punishment (purple, n=61 cells, 14 mice), or baseline sessions with no contingency (gray, n=28 cells, 10 mice) on the day of patching. Pink inset shows a zoom-in view of cells with smaller EP Sst neuron inputs. **B**, Synaptic sign index of the three conditions (reward, punishment, baseline). *Top*, synaptic sign index measures the difference between inhibitory and excitatory current amplitude normalized by total amount of input currents. Synaptic sign index closer to 1 indicates an inhibitory biased EP Sst→LHb synapses and synaptic sign index closer to -1 indicates an excitatory biased EP Sst→LHb synapses. *Middle*, average synaptic sign index of all three conditions (vertical lines) and the corresponding bootstrapped distributions used to calculate significance (semi-transparent bars). *Bottom*, cumulative distribution function of synaptic sign index of each condition.

**Fig. S12.**
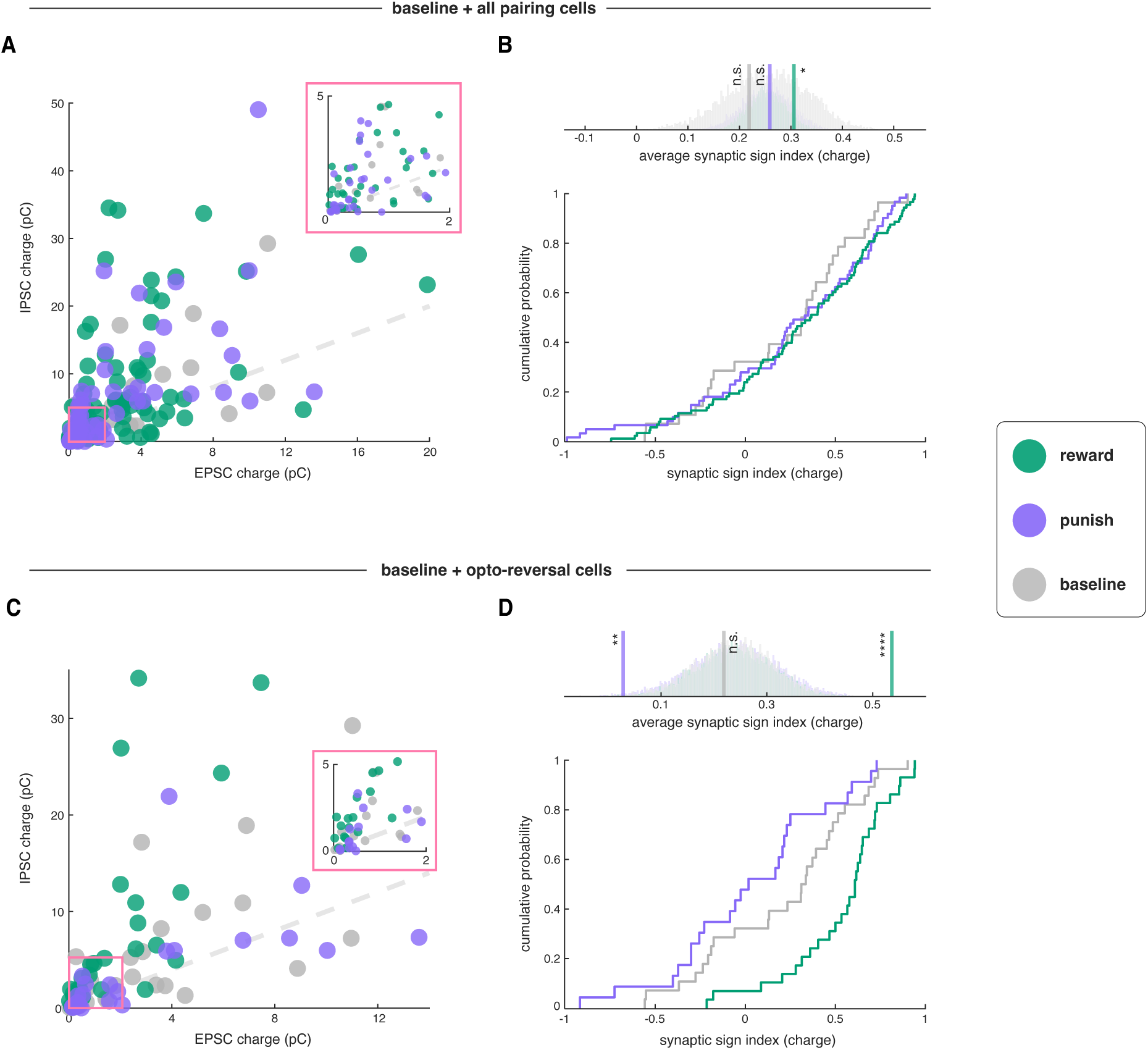
Synaptic signs of EP Sst**→**LHb synapses (classified by opto-outcome pairing direction) calculated using total charge. **A**, Scatter plot of EPSC total charge and IPSC total charge of all cells recorded. Each dot represents a cell. Cells are colored based on whether EP opto stim are paired with reward (green) or punishment (purple), or baseline sessions with no contingency (gray) on the day of patching. Pink inset shows a zoom-in view of cells with smaller EP Sst neuron inputs. **B**, Synaptic sign index (calculated by charge) of the three conditions (reward, punishment, baseline). *Top*, average synaptic sign index of all three conditions (vertical lines) and the corresponding bootstrapped distributions used to calculate significance (semi-transparent bars). *Bottom*, cumulative distribution function of synaptic sign index of each condition. **C**, Scatter plot of EPSC total charge and IPSC total charge of baseline and opto-reversal cells. Each dot represents a cell. Cells are colored the same as in **Fig. 3D**. Pink inset shows a zoom-in view of cells with smaller EP Sst neuron inputs. **D**, Synaptic sign index (calculated by charge) of the three conditions (reward, punishment, baseline). *Top*, average synaptic sign index of all three conditions (vertical lines) and the corresponding bootstrapped distributions used to calculate significance (semi-transparent bars). *Bottom*, cumulative distribution function of synaptic sign index of each condition.

**Fig. S13.**
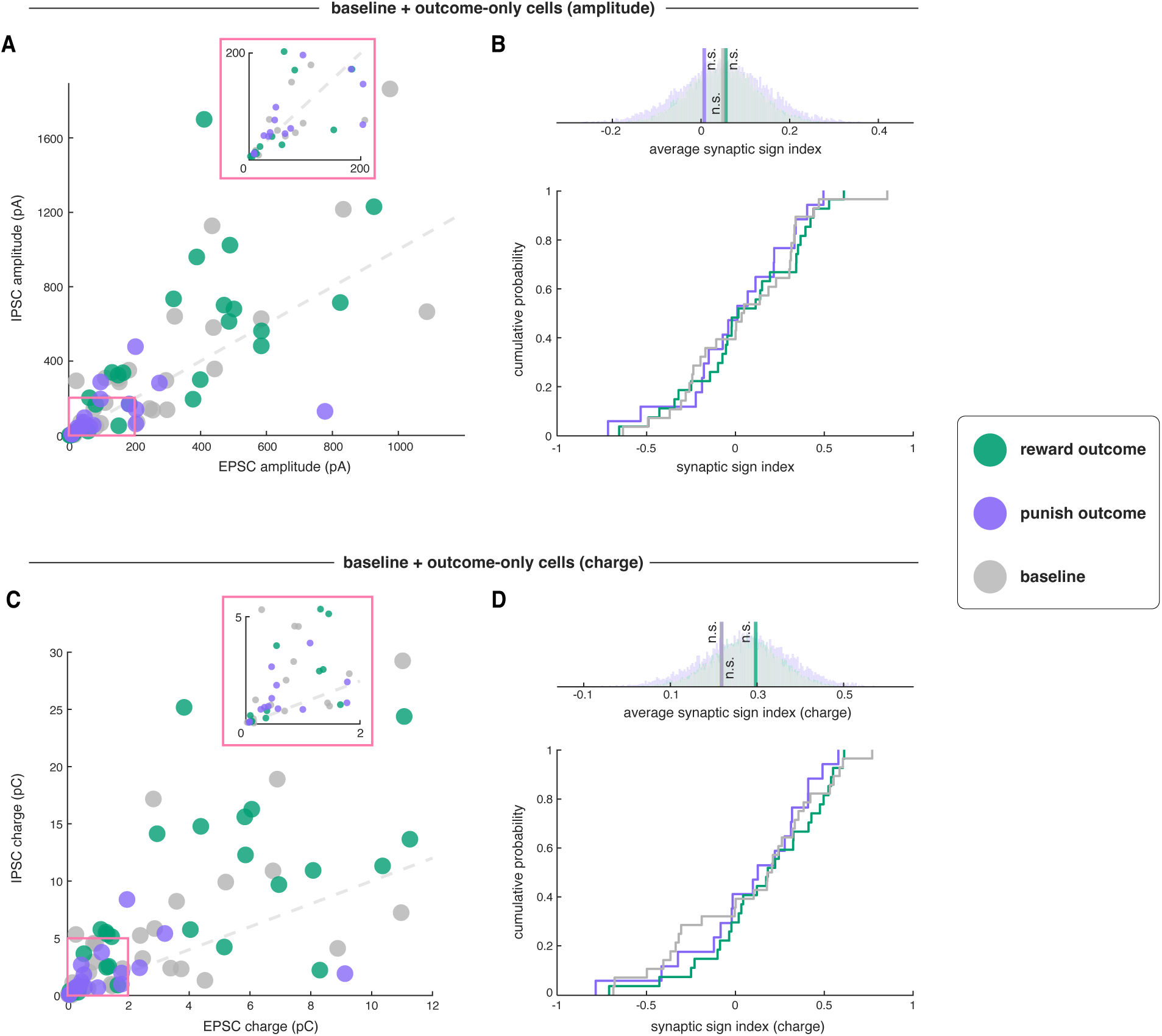
Synaptic signs of EP Sst**→**LHb synapses did not switch upon repeated exposure of reward or punishment outcome alone. **A**, Scatter plot of EPSC amplitude and IPSC amplitude of all cells recorded in animals that experienced repeated reward outcome only (water delivery) or punishment outcome only (air puffs). Each dot represents a cell. Cells are colored based on whether animal experienced reward outcome only (green, n=27 cells, 6 mice) or punishment outcome only (purple, n=17 cells, 5 mice), or baseline sessions with no contingency (gray, n=28 cells, 10 mice) on the day of patching. Pink inset shows a zoom-in view of cells with smaller EP Sst neuron inputs. **B**, Synaptic sign index of the three conditions (exposure to reward outcome, punishment outcome, or baseline). *Top*, average synaptic sign index of all three conditions (vertical lines) and the corresponding bootstrapped distributions used to calculate significance (semi-transparent bars). *Bottom*, cumulative distribution function of synaptic sign index of each condition. **C**, Scatter plot of EPSC total charge and IPSC total charge of the same cells in panel **A**. Each dot represents a cell. Cells are colored the same as panel **A**. Pink inset shows a zoom-in view of cells with smaller EP Sst neuron inputs. **D**, Synaptic sign index (calculate by charge) of the three conditions of increase, stable, or decrease DA amplitude. *Top*, average synaptic sign index of all three conditions (vertical lines) and the corresponding bootstrapped distributions used to calculate significance (semi-transparent bars). *Bottom*, cumulative distribution function of synaptic sign index of each condition.

**Fig. S14.**
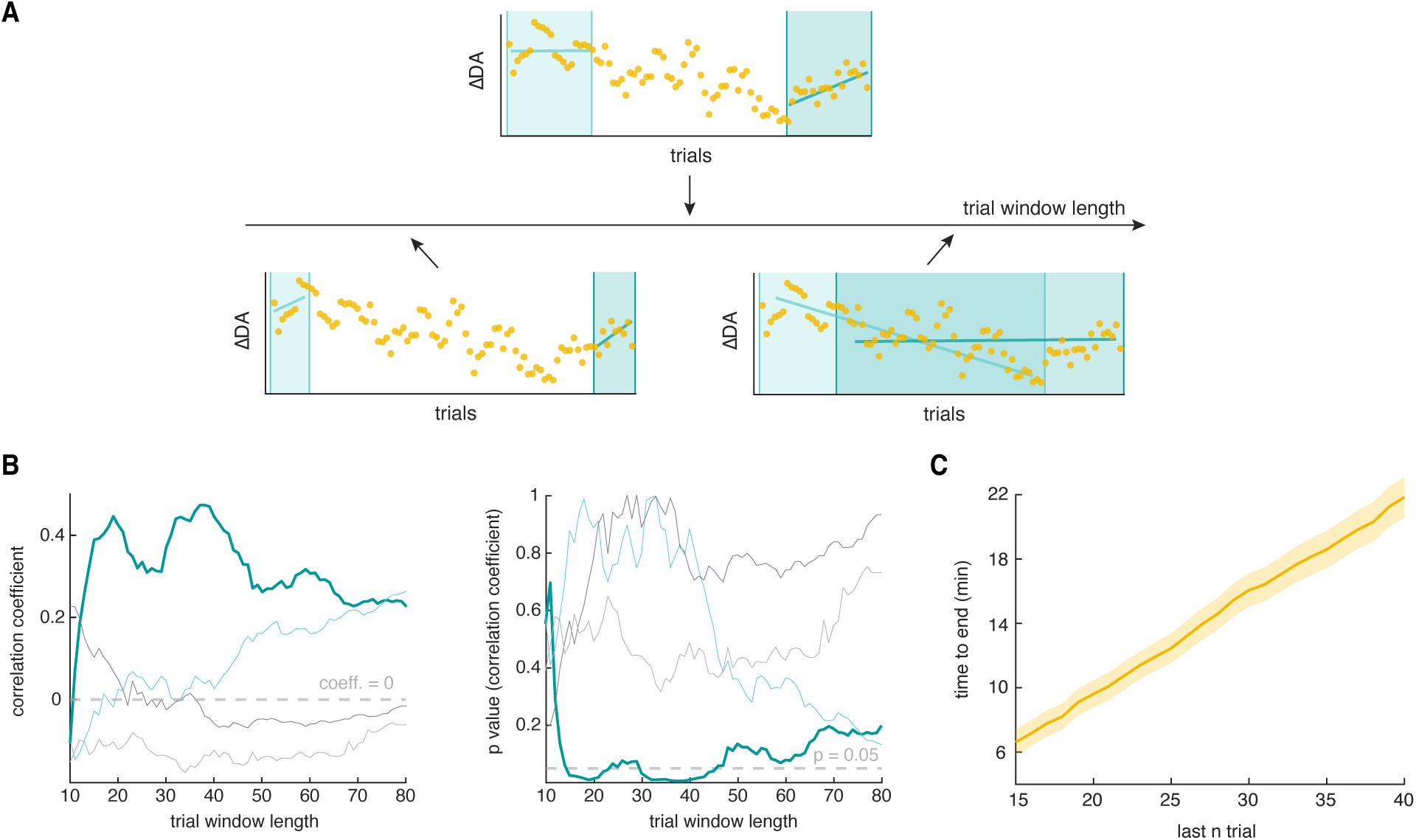
Approximate timescale of sign-switching plasticity. **A**, Schematics of analysis. For each animal, we take the latest opto-pairing sessions (i.e. on the day of *ex vivo* recording) and measure the correlation between animal sign index and DA slope during early (light green) and late (dark green) windows of all possible lengths, as well as window-averaged DA transient amplitude (light gray: aligned to the first trial; dark gray: aligned to the last trial). The trial window for early or late window varies from 10 to 80 trials (i.e. the whole session). **B**, Correlation coefficients (*left*) and corresponding p-values (*right*) of animal sign index and DA slope calculated as detailed above as a function of trial window length. Note that only DA slopes calculated during the late window showed significant correlation with animal sign index, as seen by the continuous region between last 15 to last 40 trials where DA slope significantly correlates with animal sign index. **C**, Time between the last 15 to 40 trials to the end of session. The averaged time ± s.e.m are shown.

**Fig. S15.**
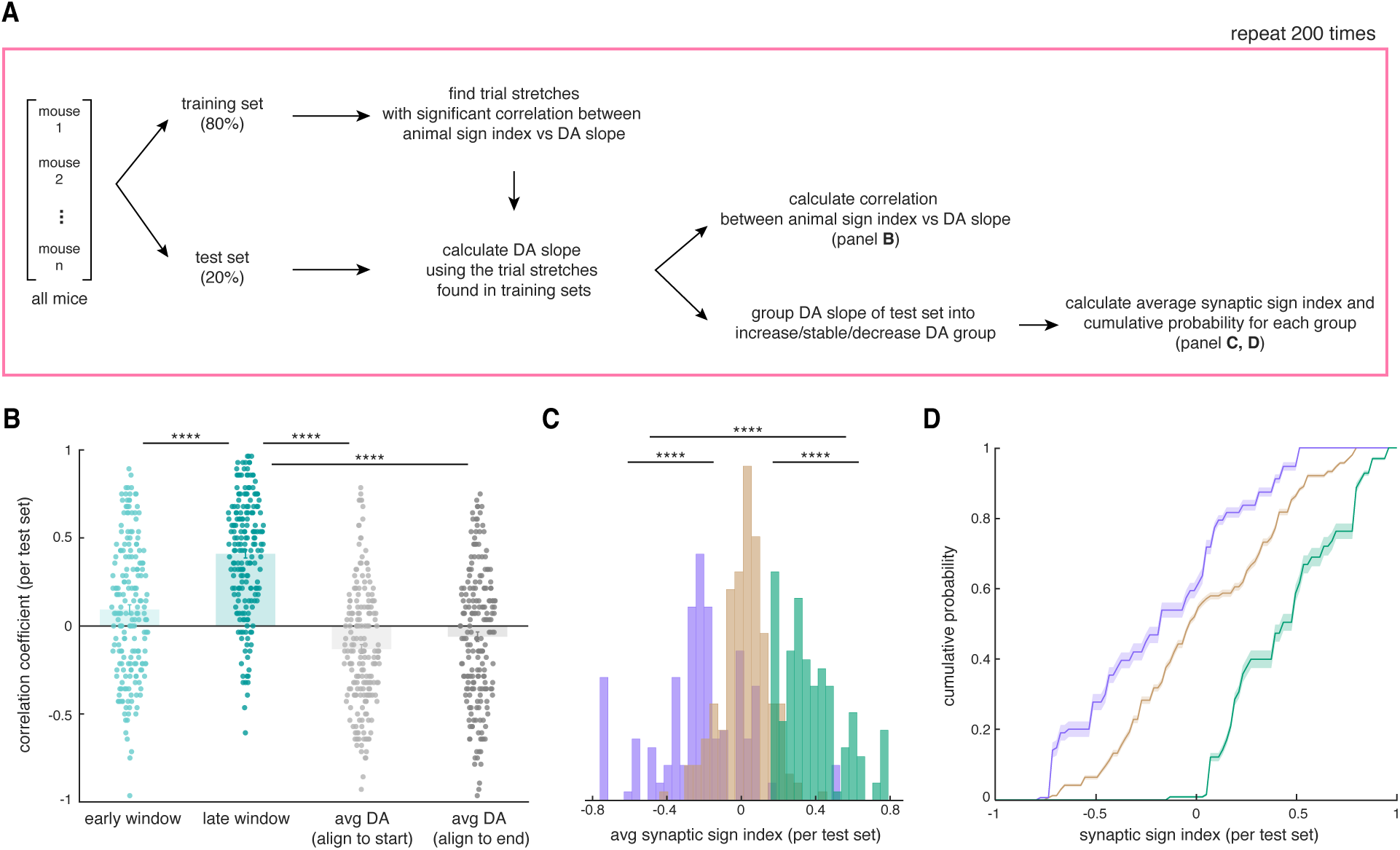
Analysis of dataset with train-test-splits. **A**, Overview of analysis pipeline. We randomly separated all animals in the dataset into training set (80% of animals) and test set (20% of animals). Using data recorded from the train animals, we identify trial stretches with significantly correlated animal sign index and DA slope of interest (early or late DA slope, average DA amplitude aligned to start or end of the session). These trial stretches found in training set are then used to calculate correlation between animal sign index and DA slope in animals from the test set. We also calculate the average synaptic sign index and cumulative probability of synaptic sign index from cells of test animals based on their respective DA slope group (increasing/stable/decreasing DA). We repeat this process for 200 times. **B**, Correlation coefficients of animal sign index and DA slope for all train-test splits calculated as detailed above. Note that only DA slopes calculated during the late window showed significant correlation with animal sign index. Each dot is the correlation coefficient for a given train-test split and the average correlation coefficient across all repeats ± s.e.m are shown. **C**, Histogram of all average synaptic sign index across all repeats from cells recorded in test animals. Cells are grouped by DA slope for each animal (green: increasing DA, brown: stable DA, purple: decreasing DA). **D**, Cumulative probability of synaptic sign index from cells recorded in test animals across all train-test split repeats. The average cumulative probability ± s.e.m are shown.

**Fig. S16.**
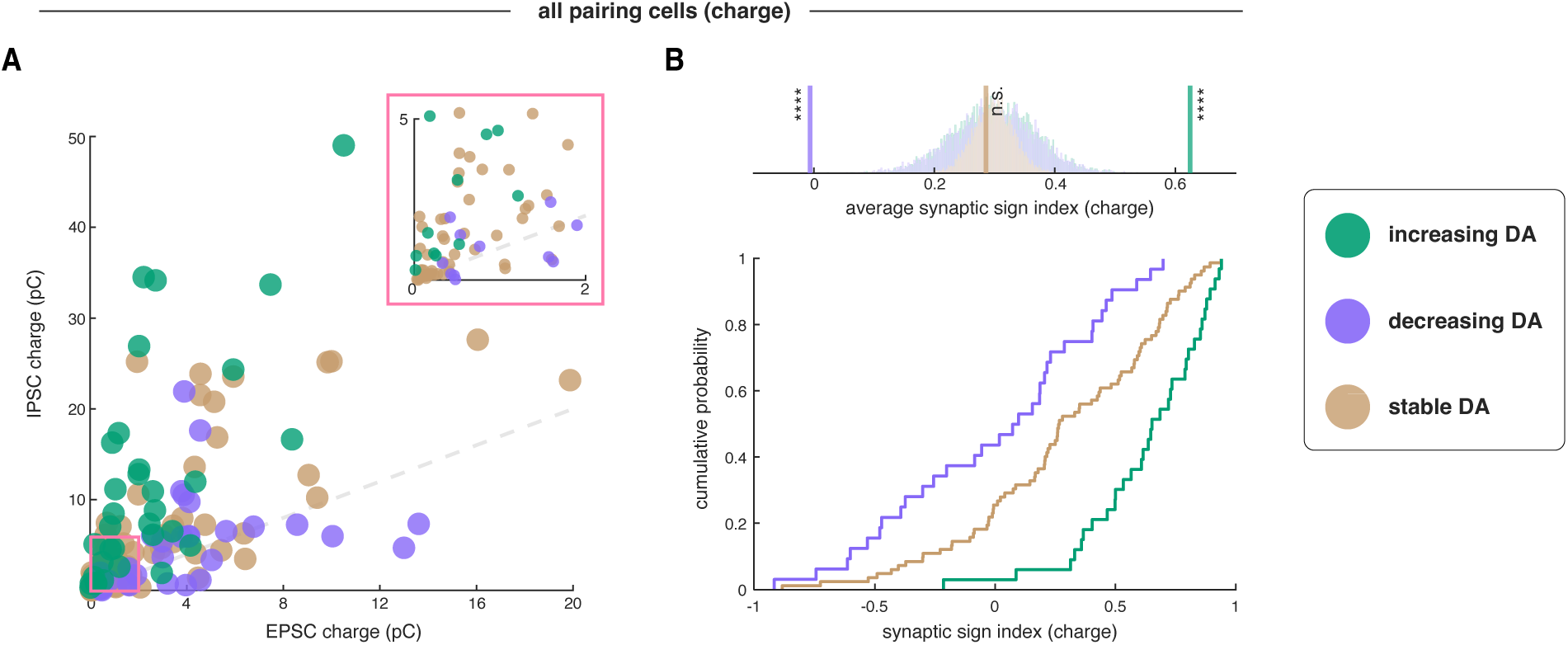
Synaptic signs of EP Sst**→**LHb synapses (classified by DA update direction) calculated using total charge. **A**, Scatter plot of EPSC total charge and IPSC total charge of all cells recorded. Each dot represents a cell. Cells are colored the same as **Fig. 4H**. Pink inset shows a zoom-in view of cells with smaller EP Sst neuron inputs. **B**, Synaptic sign index of the three conditions of increasing, stable, or decreasing DA amplitude. *Top*, average synaptic sign index of all three conditions (vertical lines) and the corresponding bootstrapped distributions used to calculate significance (semi-transparent bars). *Bottom*, cumulative distribution function of synaptic sign index of each condition.

**Fig. S17.**
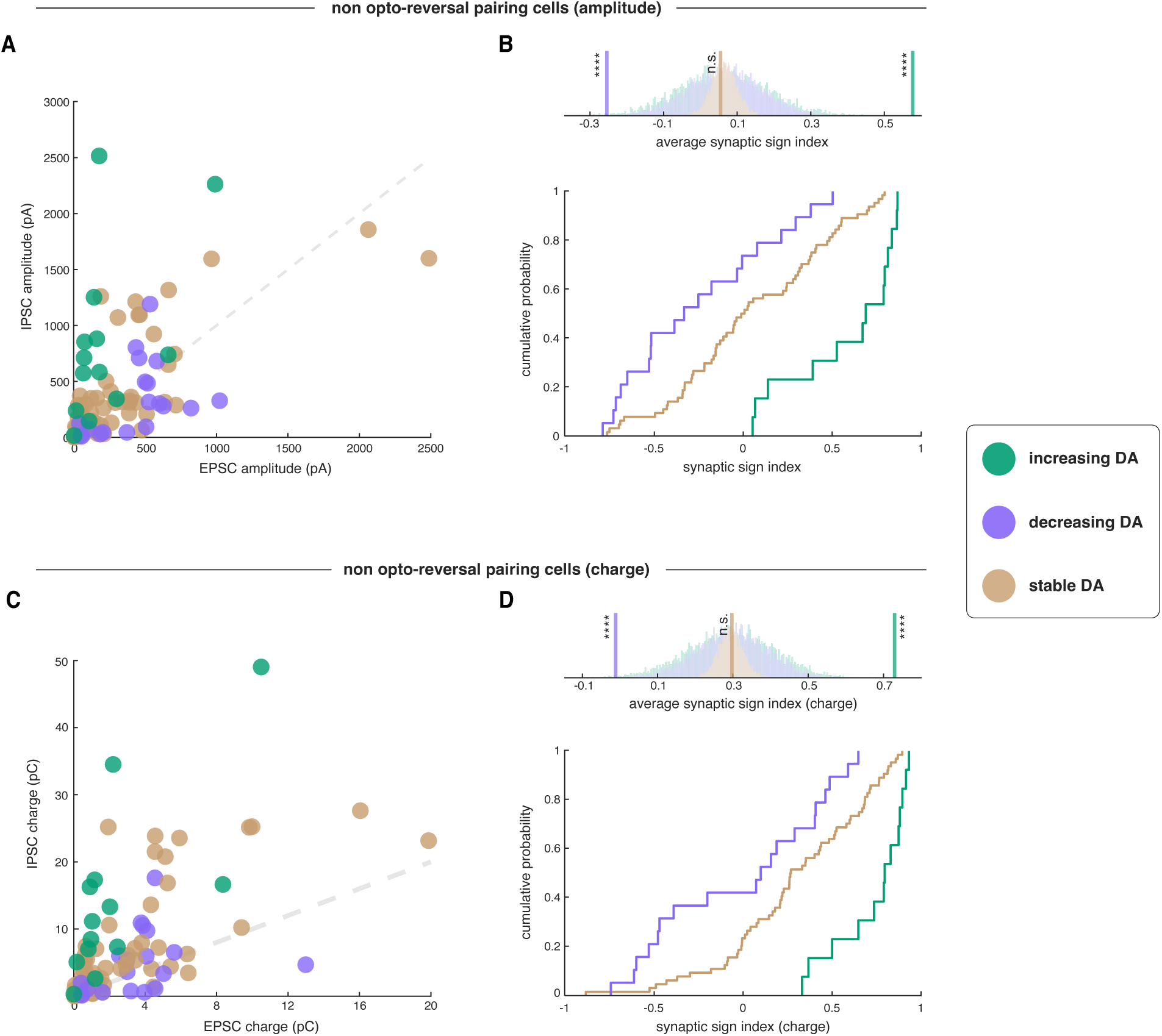
Synaptic sign switch in EP Sst**→**LHb synapses persists when excluding cells recorded during opto-outcome reversal. **A**, Scatter plot of EPSC amplitude and IPSC amplitude of all non opto-reversal pairing cells recorded. Each dot represents a cell. Cells are colored based on whether the animal experienced increasing (green, n=13 cells, 3 mice), stable (brown, n=64 cells, 15 mice), or decreasing (purple, n=19 cells, 5 mice) DA amplitude during the late window. Pink inset shows a zoom-in view of cells with smaller EP Sst neuron inputs. **B**, Synaptic sign index of the three conditions of increasing, stable, or decreasing DA amplitude. *Top*, average synaptic sign index of all three conditions (vertical lines) and the corresponding bootstrapped distributions used to calculate significance (semi-transparent bars). *Bottom*, cumulative distribution function of synaptic sign index of each condition. **C**, Scatter plot of EPSC total charge and IPSC total charge of all non opto-reversal pairing cells recorded. Each dot represents a cell. Cells are colored the same as panel **A**. Pink inset shows a zoom-in view of cells with smaller EP Sst neuron inputs. **D**, Synaptic sign index (calculate by charge) of the three conditions of increasing, stable, or decreasing DA amplitude. *Top*, average synaptic sign index of all three conditions (vertical lines) and the corresponding bootstrapped distributions used to calculate significance (semi-transparent bars). *Bottom*, cumulative distribution function of synaptic sign index of each condition.

**Movie S1. Behavior setup for *in vivo* learning and example opto+tone and opto-only trials.** Video recording of an example opto+tone trial and an example opto-only trial. Cue periods in the corresponding trials were indicated in white text in the video. Note that during optogenetic stimulation, the overhead red lamp masked the light from the EP opto stim, and there was no apparent auditory cue that could indicate the delivery of optogenetic stimulation.

Movie S2. Example opto-only trial at the start or end of reward pairing or punishment pairing sessions. Video recording of the first and last opto-only trial during reward pairing and punishment pairing. Cue periods and time of outcome delivery were indicated in white on the upper right portion of the video.

